# Bacterial and fungal components of the gut microbiome have distinct, sex-specific roles in Hawaiian *Drosophila* reproduction

**DOI:** 10.1101/2023.07.14.549088

**Authors:** Matthew J. Medeiros, Laura Seo, Aziel Macias, Donald K. Price, Joanne Y. Yew

## Abstract

Gut microbiomes provide numerous physiological benefits for host animals. The role of bacterial members of microbiomes in host physiology is well-documented. However, much less is known about the contributions and interactions of fungal members of the microbiome even though fungi are significant components of many microbiomes, including those of humans and insects. Here, we used antibacterial and antifungal drugs to manipulate the gut microbiome of a Hawaiian picture-wing *Drosophila* species, *D. grimshawi*, and identified distinct, sex-specific roles for the bacteria and fungi in microbiome community stability and reproduction. Female oogenesis, fecundity and mating drive were significantly diminished when fungal communities were suppressed. By contrast, male fecundity was more strongly affected by bacterial but not fungal populations. For males and females, suppression of both bacteria and fungi severely reduced fecundity and altered fatty acid levels and composition, implicating the importance of interkingdom interactions on reproduction and lipid metabolism. Overall, our results reveal that bacteria and fungi have distinct, sexually-dimorphic effects on host physiology and interkingdom dynamics in the gut help to maintain microbiome community stability and enhance reproduction.

## Introduction

For many animals, the gut microbiome functions as a “virtual” organ that facilitates a range of physiological functions including the provision of essential nutrients, immunoprotection, detoxification, and energy metabolism (Kucuk, 2020; Lee & Hase, 2014). In particular, the contributions of gut bacteria have been studied intensely in animal hosts from humans to insects (Kucuk, 2020). However, fungi are also significant constituents of many animal microbiomes (Li et al., 2021; Zhang et al., 2022). Far fewer beneficial roles for the mycobiome, the fungal component of the microbiome, have been identified. Notably, fungi enhance the immune system in mice (van Tilburg Bernardes et al., 2020), offer protection from pathogenic bacteria, and provide nutritional scavenging (Li et al., 2021). The contributions of bacteria-fungal interactions in animal physiology have mostly been studied in the context of pathogenesis, such as the synergistic interaction of *Candida albicans*, a widespread fungal species, and the bacterium *Pseudomonas aeruginosa* in infection and cystic fibrosis (Neely et al., 1986). Despite the prevalence of bacterial-fungal mutualisms in numerous organisms including plants (Duponnois et al., 1993; Frey-Klett et al., 2007; Khalid & Keller, 2021), nematodes (Ingham et al., 1985), insects (Aylward et al., 2014), and mammals (Hacquard et al., 2015), much less is known about the beneficial contributions of interkingdom interactions in maintaining microbiome community structure and supporting host fitness.

The Hawaiian *Drosophila* clade is a powerful model for delineating the contribution of both bacterial and fungal kingdoms to host physiology. The subclade of picture wing *Drosophila* (PWDs) are endemic to the Hawaiian Islands and are notable for their relatively recent isolation and rapid speciation, with the majority of species having originated less than 3 mya (Magnacca & Price, 2015). As with other *Drosophila* groups, PWDs have a mutualistic relationship with yeast (Biedermann & Vega, 2020). However, in contrast to continental drosophilids and lab raised flies, Hawaiian PWD harbor a rich diversity of both fungi and bacteria in their gut despite having been raised for multiple generations in the lab (this study; (Chandler et al., 2012)). Importantly, a number of species are considered to be specialist feeders who appear to rely on particular sets of fungi, mostly from the genus *Saccharomyces*, as a source of their nutrition and identifier of host plants (O’Connor et al., 2014; Ort et al., 2012). The intricate co-evolution and co-dependency of PWDs and fungi provide an exceptional opportunity for understanding how bacterial and fungal components of the microbiome support host adaptation and fitness.

We used *D. grimshawi*, a member of the PWD clade of flies, to dissect how bacteria, fungi, and their interactions modulate host physiology and microbiome stability. We manipulated the abundance of each kingdom using antimicrobial drugs and measured the ensuing effects on reproduction, mating behavior, cuticular lipids, and fatty acid levels. Our findings reveal that the bacterial and fungal components of the gut microbiome play different roles in reproduction for each sex. Female fecundity and mating drive are highly dependent on gut fungal communities whereas male fecundity is influenced more by gut bacteria. Additionally, alterations in reproductive function are accompanied by sex-specific changes in cuticular lipid and fatty acid levels. Notably, the suppression of both microbial kingdoms results in an almost complete suppression of fecundity for both females and males, an outcome that can be partly rescued through fecal microbiome transplants.

## Methods

### Drosophila husbandry

The flies used for this study derive from an approximately 50 year old lab stock of *D. grimshawi*, a generalist species native to Maui Nui (recorded from Maui, Molokaʻi, Lanaʻi). Flies were reared on a standard diet consisting of 90 mL water, 1 g agar, 24 g Gerber baby banana food, 3 g powder mix (made by blending equal parts wheat germ, textured soy protein, and Kellog’s Special K cereal), 375 μL proprionic acid (Avantor; Radnor, PA), and 375 μL 100% non-denatured ethanol (Decon Labs Inc., King of Prussia, PA). For experimental assays. *D. grimshawi* were removed from rearing jars within one week of emergence, sexed, and placed into gallon-sized glass jars lined along the bottom with moist sand.Fly cultures and all experimental treatments and assays were maintained in an incubator held at 18 °C, 60% relative humidity with a 12 hr/ 12 hr day/ night cycle.

### Antimicrobial treatment

Flies were separated by sex within 7 days of eclosion and raised on one of the following types of food for 21 d: standard diet (control), standard diet supplemented with coconut oil (COil), antifungal treatment (AF), antibacterial treatment (AB), and antibacterial and antifungal treatment (AB+AF). The AB food consisted of standard media containing 200 mg/ mL of ampicillin and kanamycin (VWR Life Science; both dissolved in water), 50 mg/ mL of tetracycline (EMD Millipore Corp.; dissolved in 70% ethanol), and 300 mg/ mL erythromycin (Acros Organics; dissolved in 100% ethanol). The AF food contained 1.25 mM captan (N-trichloromethylmercapto-4-cyclohexene-l,2-dicarboximide; Sigma-Aldrich) with coconut oil (300 μL/ L standard media; Kirkland brand) used to suspend the captan. The AF+AB treatment included captan, coconut oil, and each of the antibacterial drugs used at the concentrations described above. The COil food contained coconut oil (300 μL/ L) and served as the control condition for experiments with AF or AB+AF treatments.

### Plating Drosophila tissue

Single flies were cold-anesthetized, rinsed twice in 2.5% bleach solution, and homogenized in 100 μL phosphate buffered saline (pH 7.4) for 10 s using a hand-held homogenizer. Ten μL each of dilutions (10^-1^, 10^-2^, 10-^3^) of each homogenate were plated onto YPD media with 50 μg/ mL ampicillin, kanamycin, and tetracycline, and 15 μg/ mL erythromycin (to asses fungal growth) and MRS with 0.38 mg/ mL captan (to assess bacterial growth). Plates were incubated at 30 °C for 2 days and microbial colony forming units (CFUs) counted.

### High throughput sequencing

Flies were surface sterilized with 2 washes in 95% EtOH followed by 2 washes in sterile water. Six to eight flies were prepared for each condition, with equal numbers of males and females. Individual flies were homogenized in ATL buffer from PowerMag Bead Solution (Qiagen) with 1.4 mm ceramic beads (Qiagen; MD, USA) using a bead mill homogenizer (Bead Ruptor Elite, Omni, Inc; GA, USA) and extended vortexing for 45 min at 4 °C. The homogenate was treated overnight with proteinase K (2 mg/ mL) at 56 °C and DNA was extracted using the MagAttract PowerSoil DNA EP Kit (Qiagen) according to manufacturer’s instructions. Bacterial diversity was characterized by PCR amplification of the 16S rRNA gene with primers to the V3-V4 region (515F: GTGYCAGCMGCCGCGGTAA; 806R: GGACTACNVGGGTWTCTAAT) (Parada et al., 2016). Fungal diversity was characterized using primers to the internal transcribed spacer (ITS1f: CTTGGTCATTTAGAGGAAGTAA; ITS2: GCTGCGTTCTTCATCGATGC) (White et al., 1990). The primers contain a 12-base pair Golay-indexed code for demultiplexing.

PCRs were performed with the KAPA3G Plant kit (Sigma Aldrich, MO, USA) using the following conditions: 95 °C for 3 min, followed by 35 cycles of 95 °C for 20 seconds, 50 °C for 15 seconds, 72 °C for 30 seconds, and a final extension for 72 °C for 3 min. The PCR products were cleaned and normalized with the Just-a-plate kit (Charm Biotech, MO, USA). High throughput sequencing (HTS) was performed with Illumina MiSeq and 250 bp paired-end kits (Illumina, Inc., CA, USA).

### High throughput sequencing data analysis

Post-processing of HTS data (read filtering, denoising, and merging) was performed using the “MetaFlow|mics’’ microbial 16S pipeline for bacteria and the Fungal ITS pipeline for fungi (Arisdakessian et al., 2020). Reads shorter than 20 bp and samples with fewer than 5,000 reads were discarded. Paired reads are merged if the overlap is at least 20 bp with a maximum 1 bp mismatch. The contigs generated by DADA2 (Callahan et al., 2016) were processed using MOTHUR (Schloss et al., 2009) and initially aligned and annotated using the SILVA v138 database (Quast et al., 2013). We chose a 97% sequence similarity cutoff for determination of OTUs as we were attempting to confirm the efficacy of the antimicrobial drugs rather than investigate effects to specific strains of microbes. Sequences that were unassigned by the pipeline were identified by manual searches in NCBI BLAST, UNITE (Nilsson et al., 2019), and MycoBank (Robert et al., 2013) using a >95% sequence similarity cutoff. The ITS data were normalized in R using the DESeq2 package (Love et al., 2014). Analyses were performed after clustering at the genus level using R version 4.2.1, and the phyloseq package (McMurdie & Holmes, 2013). Alpha diversity was measured using Chao1 and Shannon diversity metrics and a Wilcoxon statistical test. Beta diversity was calculated using non-metric multi-dimensional scaling (NMDS using binary Jaccard distances) and an analysis of similarities (ANOSIM) test. Relative abundance charts were constructed after grouping flies from the same treatment. Univariate multiple testing with an F-test, was used to test for significant differences in microbial taxa (phyloseq). Data from female and male flies were pooled.

### Fecundity

Following 3 weeks of antimicrobial or control treatment, a single male and single female were placed in a standard 8-dram fresh control food vial for 48 hrs, after which males were removed. Food for egg laying trials contained one drop of blue food coloring (Spice Supreme, Bayonne, NJ) per batch to increase the visibility of eggs. Females were transferred to new food vials twice per week and outgoing food vials were checked for eggs for the lifespan of the female. Vials containing eggs were checked for larvae twice a week.

For ovary development studies, virgin female flies were fed AB+AF food for 7, 14 or 21 days. Ovaries were dissected on day 21 and the number of mature eggs were counted under 10X magnification.

### Mating behavior

Flies were maintained on experimental or control diets for 3 weeks. A single virgin male and single cirgin female were placed in a polystyrene Petri dish (60 x 15 mm) containing 0.5 mL of control food and a small piece of moistened filter paper. Each experimental diet was tested in parallel with its respective control. Each trial consisted of 36 dishes with 9 replicates of each pairwise mating combination: control male + control female, control male + treatment female, treatment male + control female, and treatment male + treatment female. Mating behavior was monitored with Raspberry Pi computers outfitted with a camera (CanaKit Raspberry Pi 4 8GB computers with Longruner 1080p HD Webcam 5MP OV5647 IR-Cut Video cameras), with the location of Petri dishes randomized. Images of the flies were taken every 60 or 90 s for a duration of 48 h and subsequently analyzed manually for copulation events, defined as the male positioned directly behind the female with both female wings expanded.

### Ovary dissections

Flies were anesthetized on ice and ovaries were dissected in PBS after 21 days of treatment. Mature eggs were manually counted under 10x magnification. For ovaries used in images, the tissue was fixed in 4% paraformaldehyde for 20 min. and washed thrice with PBS + 0.1% TritonX-100 (PBST). Ovaries were dyed with Hoescht dye (1 μg/ mL) for 10 min, washed in PBS for 10 min., and mounted on slides with SlowFade Diamond Antifade mountant (ThermoFisher Scientific, Waltham, MA). Images were obtained by epifluorescence microscopy on an Olympus BX51 microscope equipped with a Leica DFC 7000 T color digital camera.

### Fecal transplant rescue

For fecal transplant experiments, virgin male and female flies were placed on AB+AF for 7 d before switching to control food that was inoculated with 15 μL of a fecal transfer wash (in PBS). The wash was prepared from flies of the same age fed on a control diet. The fecal transplant was generated by washing the sides of the vial with 200 μL sterile PBS (care was taken not to touch the surface of the food). The droplet was collected and divided into two aliquots. One aliquot was heat inactivated by placing the wash in an 80 °C oven for 10 min. Both active and inactive washes were added to fresh food vials and allowed to evaporate prior to adding flies. At 21 d, female flies were dissected and mature eggs were counted. Testes from male flies at 21 d were dissected for fatty acid analysis (see below).

### Cuticular hydrocarbon extraction

Flies were cold anesthetized at 4 °C, placed in glass vials, and covered with 240 μL of hexane spiked with 10 ug/ mL hexacosane for 10 min at RT. Next, 200 μL of solvent was removed, added to a fresh vial and evaporated under a gentle stream of N2. Samples were frozen at -20°C until analysis by gas chromatography mass spectrometry (GCMS).

### Fatty acid extraction

Flies were stored at -80 °C until analysis. Each replicate consisted of 3 flies. Samples were homogenized in 100 μL MilliQ water with 4 ceramic beads (1.4 mm diam.; Millipore) using an Omni BeadRuptor for 30 sec at 6 m/s. Twenty μL of the homogenate was removed for protein quantification using a bicinchoninic acid assay (BCA) kit (ThermoFisher). Lipids were extracted with chloroform: MeOH (2:1 v:v) spiked with 10 μg/ mL pentadecanoic acid as an internal standard. After agitation at 4 °C for 3 hours, the lower chloroform phase was removed and placed in a clean vial. The homogenate was re-extracted twice with chloroform. Solvent from the pooled extracts was evaporated under N2 and stored at -20 °C until esterification. Samples were esterified with 200 µL of 0.5 N methanolic HCl (Sigma Aldrich, St. Louis, MO) at 65 °C for 1.5 hours with occasional vortexing. Following solvent evaporation, fatty acids were reconstituted in 100 µL hexane prior to GCMS analysis.

### Gas chromatography mass spectrometry (GCMS)

GCMS analysis was performed on a 7820A GC system equipped with a 5975 Mass Selective Detector (Agilent Technologies, Inc., Santa Clara, CA, USA) and a HP-5ms column ((5%-Phenyl)-methylpolysiloxane, 30 m length, 250 μm ID, 0.25 μm film thickness; Agilent Technologies, Inc.). Electron ionization (EI) energy was set at 70 eV. One microliter of the sample was injected in splitless mode and analyzed with helium flow at 1 mL/ min. The following parameters were used for fatty acids (FA): the oven was initially set at 50 °C for 2 min, increased to 90 °C at a rate of 20 °C/min and held at 90 °C for 1 min, increased to 280 °C at a rate of 5 °C/min and held at 280 °C for 2 min. For cuticular hydrocarbon (CHC) analysis, the oven was initially set at 40 °C for 3 min, increased to 200 °C at a rate of 35 °C/ min, increased to 280 °C at a rate of 20 °C/ min, and held at 280 °C for 15 min. The MS was set to detect from m/z 33 to 500. Data were analyzed using MSD ChemStation (Agilent Technologies, Inc.).

### Fatty acid and cuticular hydrocarbon analysis

Individual FA species were identified on the basis of retention time and characteristic fragmentation patterns compared to that of standards in the National Institute of Standards and Technology database (NIST 98). The area under the peak of each FA was integrated, normalized to the area of the spiked standard, and summed to determine total FA levels. To analyze changes in length between experimental and control conditions, the intensity of individual FA signals was normalized to the total peak area of all FA peaks, generating proportional values for each FA. To eliminate multi-collinearity, logcontrasts were calculated for each FA peak using the following formula:

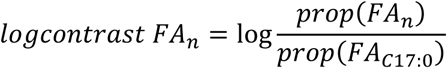

where *n* represents one of the fatty acid species. A minor saturated C17 fatty acid component was used as the divisor. Relative FA abundances were pooled according to carbon chain length (C12, C14, C16, C18, C20) or to double bond number (0, 1, or 2) and analyzed by simple linear regression.

For CHC analysis, the abundance of each CHC was quantified by normalizing the area under each CHC peak to the area of the hexacosane signal using home built peak selection software (personal correspondence, Dr. Scott Pletcher, Univ. of Michigan). To calculate total CHC levels, the normalized peak area for each CHC species was summed.

## Results

### Manipulation of gut bacteria and fungi communities

To elucidate the separate contributions of bacterial and fungal microbiome components to host physiology, we first fed antimicrobial drugs to female and male *D. grimshawi* for 21 days to suppress the growth of bacteria, fungi, or both communities. Bacteria were targeted using a cocktail of kanamycin, ampicillin, tetracycline, and erythromycin. Fungi were suppressed with the broad spectrum fungicide captan, known to inhibit ascomycete fungi, epiphytic and wine yeasts including *Saccharomyces cerevisiae (Magoye et al., 2020; Scariot FJ, 2016)*. We initially tested a second common fungicide, benomyl (1-(Butylcarbamoyl)-1*H*-1,3-benzimidazol-2-yl methylcarbamate) (Summerbell, 1993) but found it to be lethal for flies after 1-2 weeks of administration. To evaluate the effect of the drug treatments on microbiome abundance and composition, we plated whole fly homogenates on MRS media supplemented with captan (to select for bacterial growth) and YPD media supplemented with antibiotics (to select for fungal growth). Homogenates of flies treated with antifungal or antibacterial drugs resulted in significantly fewer CFUs compared to control flies (n=6 per condition), indicating that 3 weeks of treatment were sufficient to significantly suppress microbial growth in the gut and that the antibacterials and captan are effective and selective inhibitors of *Drosophila* gut bacteria and fungi, respectively (**Fig. 1**).

**Figure 1.**
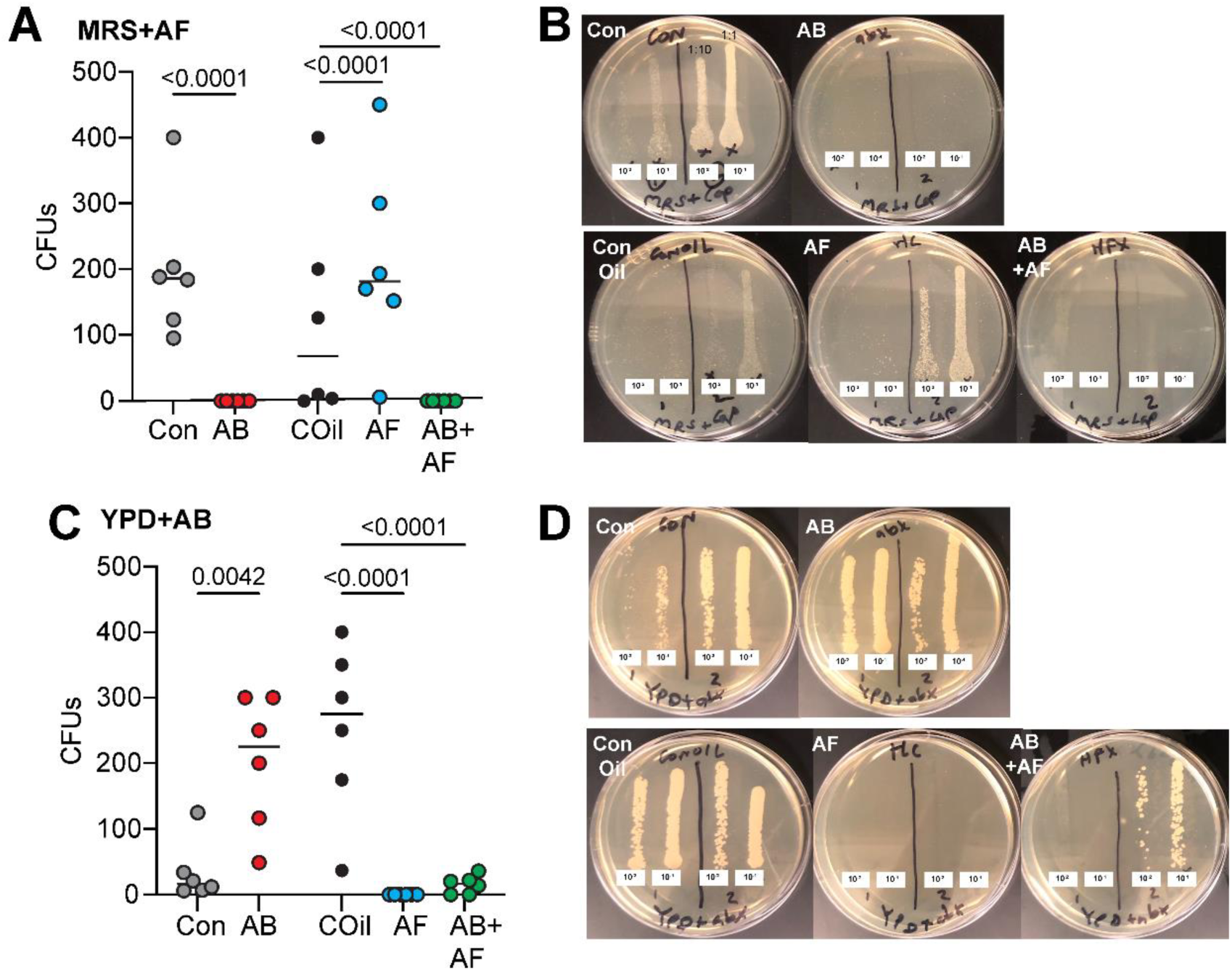
Efficacy of anti-microbial treatments. **A.** Antibacterial treatment (AB) eliminated almost all bacterial growth. Significantly fewer colony forming units (CFUs) appeaed compared to controls (Con). Treatment with the antifungal captan (AF) led to an increase in bacterial CFUs relative to the respective coconut oil-containing diet (COil). The combined AB+AF treatment suppressed almost all bacterial growth. Individual circles represent CFU counts from single fly homogenate; line indicates median. The p values were determined by a general linear model with a Poisson regression, n=6 for all treatments. **B.** Representative MRS+AF plates containing homogenate from single flies plated at two concentrations (10^-1^, 10-^2^), with 2 flies per plate. **C.** AF treatment suppressed fungal CFU growth compared to COil conditions. AB treatment led to an increase in fungal CFUs relative to the control diet. The AF and combined AB+AF treatments significantly reduced fungal growth. Individual circles represent CFU counts from single fly homogenate; line indicates median. The p values were determined by a general linear model with a Poisson regression; n=6 for all treatments. **D.** Representative YPD+AB plates containing homogenate from single flies plated at two concentrations (10^-1^, 10-^2^), with 2 flies per plate.

### Changes in microbial community composition

To determine how microbial community structure changes after treatment and whether each kingdom influences the others’ composition, we performed high throughput sequencing (HTS) of 16S rRNA and ITS amplicons from individual flies and analyzed two metrics of community composition, alpha diversity (taxonomic richness within a community) and beta diversity (similarity between communities). The most abundant bacterial genus present in control *D. grimshawi* was *Gluconobacter*, a known commensal of wild *Drosophila* (**Fig. 2**) (Staubach et al., 2013). Antibacterial (AB) treatment substantially reduced bacterial alpha diversity (**Supp. Fig. 1A**) but not fungal diversity (**Supp. Fig. 1D**). In addition, both AB and combined antibacterial and antifungal (AB+AF) treatments resulted in a significant increase of *Providencia* (**Fig. 2A, C**). By contrast, suppressing fungal growth induced a slight but non-significant increase of bacterial and fungal diversity (**Fig. 2B; Supp. Fig. 1**). Feeding flies both antibacterial and antifungal agents had little impact on 16S alpha diversity (**Supp. Fig. 1**), indicating that the simultaneous suppression of both microbial kingdoms may have negated each kingdom’s impact on the overall community architecture.

**Figure 2.**
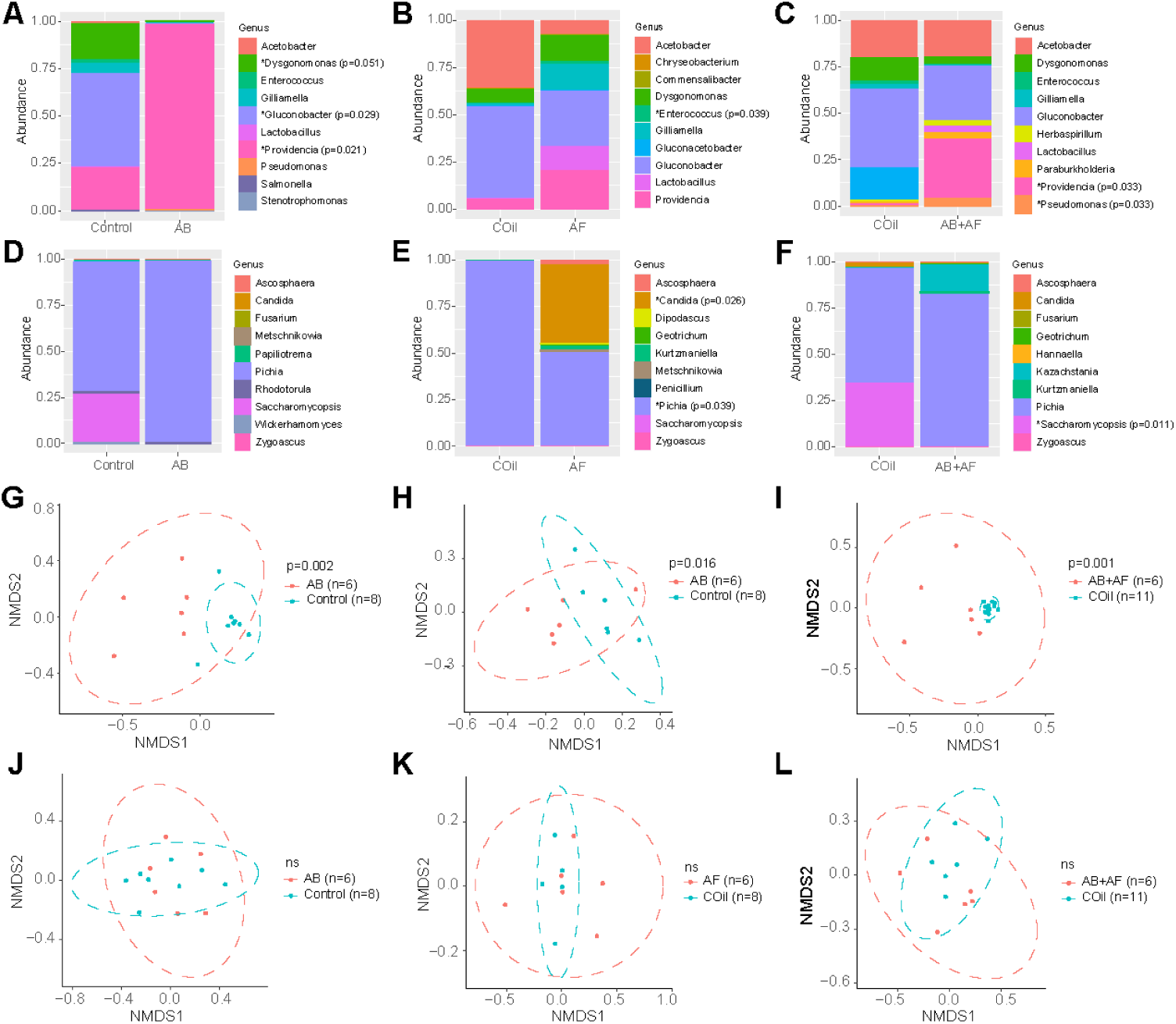
Microbiome profiles of control and experimental flies. **A – C.** Scaled relative abundance plots for the 10 most abundant bacterial genera following antibacterial (AB), antifungal (AF) or antibacterial and antifungal (AB+AF) treatment and the respective controls (Con or COil). Each column represents the average abundances of 6-8 individuals based on high throughput 16S rRNA gene amplicon sequencing. The p values were determined using univariate multiple testing with an F test; *: abundance is significantly different between control and treatment conditions. **D – F.** Scaled relative abundance plots for the 10 most abundant fungal genera following AB, AF, or AB+AF treatment or the respective controls (Con or COil). Each column represents the average abundances of 6-8 individuals based on high throughput ITS gene amplicon sequencing. The p values were determined using univariate multiple testing with an F test; *: abundance is significantly different between control and treatment conditions. **G – I.** Non-multidimensional scaling plots (NMDS; based on Jaccard distances) of OTUs (grouped at 97% similarity level) reveal distinct bacterial communities following AB, AF, or AB+AF treatment (ANOSIM). Ellipses represent significance at 0.05 confidence. **J – L.** Non-multidimensional scaling plots (NMDS; based on Jaccard distances) of OTUs (grouped at 97% similarity level) show little change in the fungal composition following AB, AF, or AB+AF treatment (all treatments p>0.05, ANOSIM). Ellipses represent significance at 0.05 confidence.

With respect to the fungal profile, *Pichia* was the most common genus in both control and treated flies (**Fig. 2D-F**). Antibacterial treatment had no significant impact on fungal abundance (**Fig. 2D**). Captan diminished the relative abundances of *Candida* and *Pichia* genera (**Fig. 2E**). In addition, *Saccharomycopsis* levels also decreased in response to AB+AF conditions (**Fig. 2F**).

Next, we examined how microbiome compositions change in response to each antimicrobial treatment. Suppressing bacterial or fungal growth either separately or concurrently altered the bacterial community composition compared to the respective controls (**Fig. 2G-L**). The combined AB+AF treatment also changed bacterial composition in a manner that was distinct from either drug alone (**Fig 2I**). In contrast to the bacterial response, the composition of the mycobiome was not significantly altered by any of the drug treatments (**Fig. 2J-L**). Interestingly, captan treatment led to sex-specific differences in microbiome composition (**Supp. Fig. 2**), a shift that was mostly driven by a decrease of *Acetobacter* and *Lactobacillus* in females compared to males. In addition, abundance of the yeast genera *Pichia* dropped precipitously in captan-treated females compared to males (**Supp. Fig. 2C**). In control flies, no differences were found between the microbiome composition of males and females.

Taken together, manipulations of each microbial kingdom separately and together reveal that the gut fungal community is more resilient to compositional changes, whereas bacterial community stability appears to be partly dependent on the composition of the fungal microbiome. Our findings also indicate that males and females respond differently to antifungal treatment.

### Distinct roles of bacteria and fungi in host reproduction

After confirming that the antimicrobial treatments were effective in suppressing the targeted microbe population and were capable of inducing significant changes in community structure, we next characterized the impact of each kingdom on reproduction and related physiological features.

#### Fecundity

Previous studies of gnotobiotic and axenic flies established that gut bacteria support host fecundity and fertility (Elgart et al., 2016; Gould et al., 2018). To determine whether *D. grimshawi* exhibit a similar dependency on gut microbes, we first measured fecundity in flies after treatment with both antibacterial and antifungal drugs. Normally, lab-raised *D. grimshawi* females begin to develop ovaries between 7-14 days post-eclosion and reach full sexual maturity with ovaries containing Stage 13/ 14 eggs between 14-21 days post-eclosion (this study; (Craddock & Boake, 1992) (**Fig. 3**). When the fly microbiome is suppressed during the first week of adult development, no mature eggs develop (**Fig. 3A, B**). However, egg production could be partially rescued by a transplant of active microbes from control flies or co-housing with a control fly. Of the flies inoculated with active fecal transfer, 52% developed mature eggs, with an average of 22.4 ± 5.5 (SEM) mature eggs/female compared to 75.9 ± 8.3 eggs/ female for controls (**Fig. 3C, D**). Co-housing resulted in partial rescue as well: 27% recovered ovary development, with an average of 8.9 ± 5.9 mature eggs per female. By comparison, only 7% of the flies treated with heat-inactivated fecal wash developed mature eggs (3.9 ± 2.9 eggs/ female) whilst no females recovered when co-housed with another AB+AF female. These results indicate that the gut microbiome, and specifically microbial activity, is necessary during a critical developmental window in early adulthood for ovary maturation. Moreover, based on the partial rescue provided by fecal transplant and co-housing, the loss of fecundity resulting from the selective suppression of microbial communities can be attributed to microbiome-related deficits rather than non-specific toxic effects.

**Figure 3.**
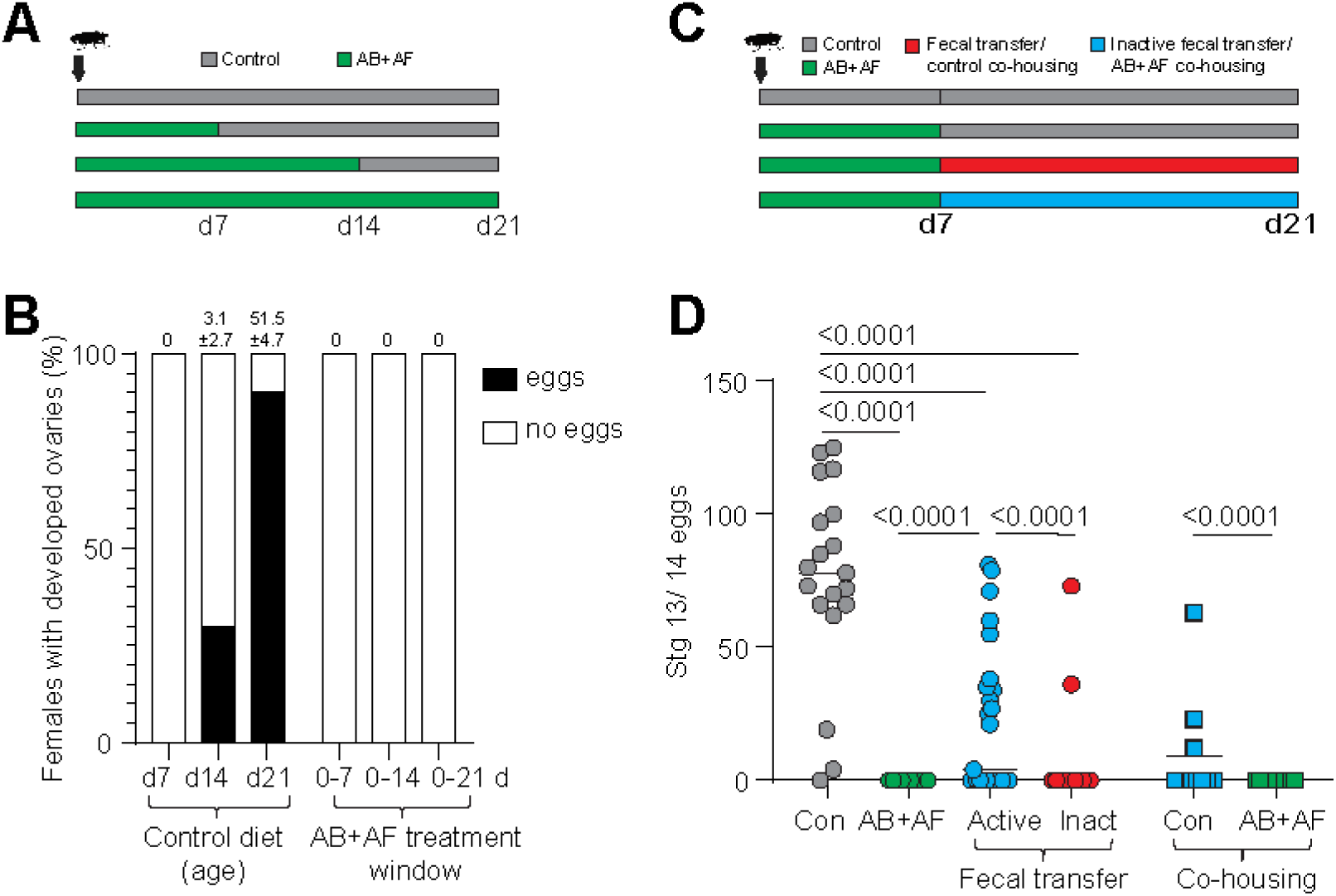
The role of the microbiome in ovary development. **A.** Feeding scheme used to test the contribution of the microbiome to oogenesis during adult development. Newly eclosed flies were continuously fed antibacterial and antifungal (AB+AF) treatment for 7, 14, or 21 d. **B.** Percentage of *D. grimshawi* with developed ovaries, defined as the presence of Stage 13/ 14 mature eggs. The majority of females fully develop ovaries between 2 - 3 weeks post-eclosion. Suppressing the microbiome during the first 3 weeks post-eclosion completely inhibits oogenesis. The average mature egg number (± standard error of mean) is indicated at the top of each bar; n=10 per condition. **C.** Feeding scheme used for antimicrobial drug administration and fecal transplant rescue. Flies are fed AB+AF treatment for 7 days then switched to control media supplemented with active fecal transplant or heat-inactivated fecal transplant for 14 days. Rescues were also performed by co-housing treatment flies with control flies or AB+AF treated flies. **D.** The AB+AF treatment suppressed egg production (Con: n=19; AB+AF: n=10). Fecal transfers from control flies (n=25) or co-housing with control flies (n=11) partially rescued oogenesis compared to heat-inactivated fecal transfer (n=28) or co-housing with AB+AF flies (n=11). Individual points represent single flies; line indicates median. The p values were determined by a general linear model with a Poisson regression.

To assess the individual roles of bacterial and fungal activity in fecundity, we first selectively suppressed each kingdom in the gut and measured ovary development in virgin females (**Fig. 4A**). Virgin females treated with antibacterial drugs developed fewer mature eggs compared to the control group (**Fig. 4B**; Con: 35.3 ± 5.1 vs AB: 19.6 ± 5.3; mean ± SEM). Antifungal treatment suppressed oogenesis to a greater degree compared to the control oil (COil) group (**Fig. 4B**; COil: 75.8 ± 8.3 vs AF: 26.2 ± 7.2). Female ovaries were minimally developed and contained no eggs for the AB+AF condition (**Fig. 4B**). Taken together, concurrent suppression of both bacteria and fungi suppressed ovary development to a greater degree than either kingdom alone.

**Figure 4.**
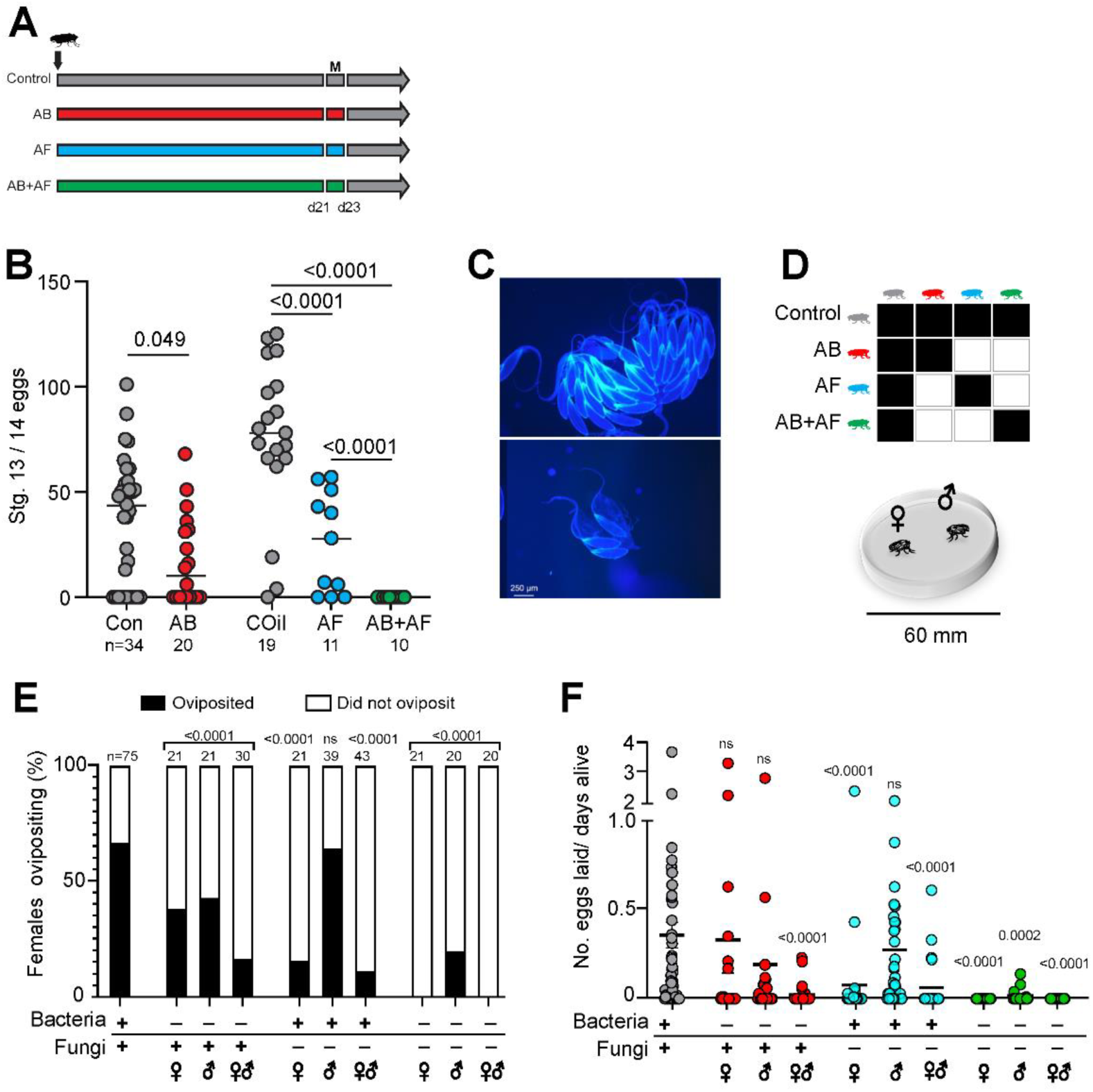
Distinct roles of bacteria and fungi in male and female fecundity. **A.** Feeding scheme to test the role of bacteria and fungi in oogenesis. Newly eclosed flies are fed antibacterial (AB), antifungal (AF), or AB+AF treatments for 21 days, mated for 2 days (M), then maintained on control food for oviposition assays. **B.** AB treatment of virgin females causes a slight decrease in the number of mature Stage 13/ 14 eggs compared to control flies. Females treated with AF or AB+AF produce significantly fewer eggs compared to the respective control. Treatment with AB+AF suppressed oogenesis to a greater extent than either drug alone. Lines indicate median. The p values were determined by a general linear model with a Poisson regression. Samples sizes are indicated beneath each treatment. **C.** Representative images of ovaries from flies fed on control oil food (top) or captan-supplemented food (bottom). **D.** Mating combinations used to test the role of bacteria and fungi in male and female fecundity. One male and one female from control, antibacterial (AB), antifungal (AF), or AB+AF treatments are placed in each courtship chamber. Flies are monitored for 48 h. **E.** AB treatment of females, males, or both sexes led to reduced fecundity. Significantly fewer AF-treated females oviposited but AF treatment had no substantial impact on male fecundity (p=0.53). Concurrent suppression of both bacteria and fungi significantly reduced male and female fecundity. Samples sizes are indicated above bars. The p values were determined by a two-tailed Fisher exact probability test. **F.** Reducing gut bacteria levels in females or males alone did not substantially change the number of eggs laid (n=21; p=0.14). However, AB female/ male dyads exhibit an additive loss of fecundity (n=30; p<.0001). Fungal suppression in females but not males significantly inhibited the number of eggs laid (n=38-39; AF females: p<0.0001). Fungal suppression in both males and females inhibits fecundity (n=43; p<0.0001)). Inhibition of both bacteria and fungi results in the loss of fecundity in both females and males compared to controls (n=20; AB+AF males: p=0.0002; AB+AF females and AB+AF dyads: p<0.0001). Lines indicate median. The p-values were determined by a Kruskall-Wallis test with Dunn’s multiple comparisons test.

We next assessed male and female fecundity in the context of mating. Female oogenesis and oviposition behavior are enhanced by mating due to the transfer of sex peptide (Liu & Kubli, 2003) and accessory gland proteins (Chen, 1996; Wolfner, 1997) from males. As such, the induction of egg laying and the number of eggs laid provide measures of both female and male fecundity. We placed single males and females in a mating chamber and measured the percentage of females that oviposited (an indicator of successful copulation) and the number of eggs laid following microbiome manipulation of males, females, or both sexes (**Fig. 4C, D**). Bacterial suppression of females or males led to a significant decrease in the percentage of females that oviposited as well as males’ ability to induce oviposition (**Fig. 4E**). However, the number of eggs laid by AB-treated females did not change compared to controls (**Fig. 4F**). By contrast, bacterial suppression in both males and females substantially decreased the number of eggs laid (**Fig. 4F**).

Suppressing fungi alone or both fungal and bacterial communities in females significantly reduced egg laying (**Fig. 4 E, F)**. Only 15.8 % of captan-fed females oviposited. Strikingly, none of the AB+AF females laid eggs **(Fig. 4E**). Considering that eliminating both microbial kingdoms resulted in a stronger effect on fecundity than suppression of either community alone, the findings indicate that bacterial-fungal interactions likely contribute to fecundity.

Male fecundity exhibited a different pattern of microbe dependency compared to females, relying substantially less on fungi. We assessed male fecundity based on the ability to induce control females to oviposit, an indicator of successful copulation (Kubli, 1992). As with females, suppressing bacterial levels had a negative impact on males’ ability to induce egg laying (**Fig. 4E, F**). However, in contrast to females, fungal suppression in males had little effect on male fecundity as indicated by two measures: the high proportion of partnered control females that oviposited and the number of eggs laid, both of which were indistinguishable from controls. When both bacteria and fungi communities were concurrently inhibited, male fecundity was profoundly suppressed (**Fig. 4E, F**). Only 20% of control females paired with AB+AF males oviposited. Of the ones that successfully mated, the egg laying rate was reduced to 0.01 eggs/ day, compared with 0.4 eggs/ day for controls. As with females, the effect of reducing the presence of both kingdoms in the gut was stronger than manipulation of either microbe community alone. In summary, the microbiome and in particular, the interactions between bacteria and fungi are essential for male and female fecundity and oogenesis. Notably, female reproduction depends more on the fungal community whereas male fecundity relies more on bacterial activity.

#### Mating

Our observations that AF and AF+AB treatments resulted in females laying fewer eggs can be partly explained by a loss of fecundity. However, it may also be the case that fewer of the treated flies copulated. To address whether the gut microbiome influences mating decisions, we counted the frequency of copulation events in paired males and females (**Fig. 5A**). Control dyads mated 73% of the time. Both males and females on AB diets exhibited little change in mating frequency compared to control flies. However, significantly fewer females laid eggs despite mating (p<0.0001, Fisher Exact Probability test), indicating that with AB treatments, fewer copulations were successful at inducing oviposition behavior (**Fig. 4E**). As such, the reduction in male and female fecundity following AB treatment is primarily due to a change in reproductive physiology rather than mating drive.

**Figure 5.**
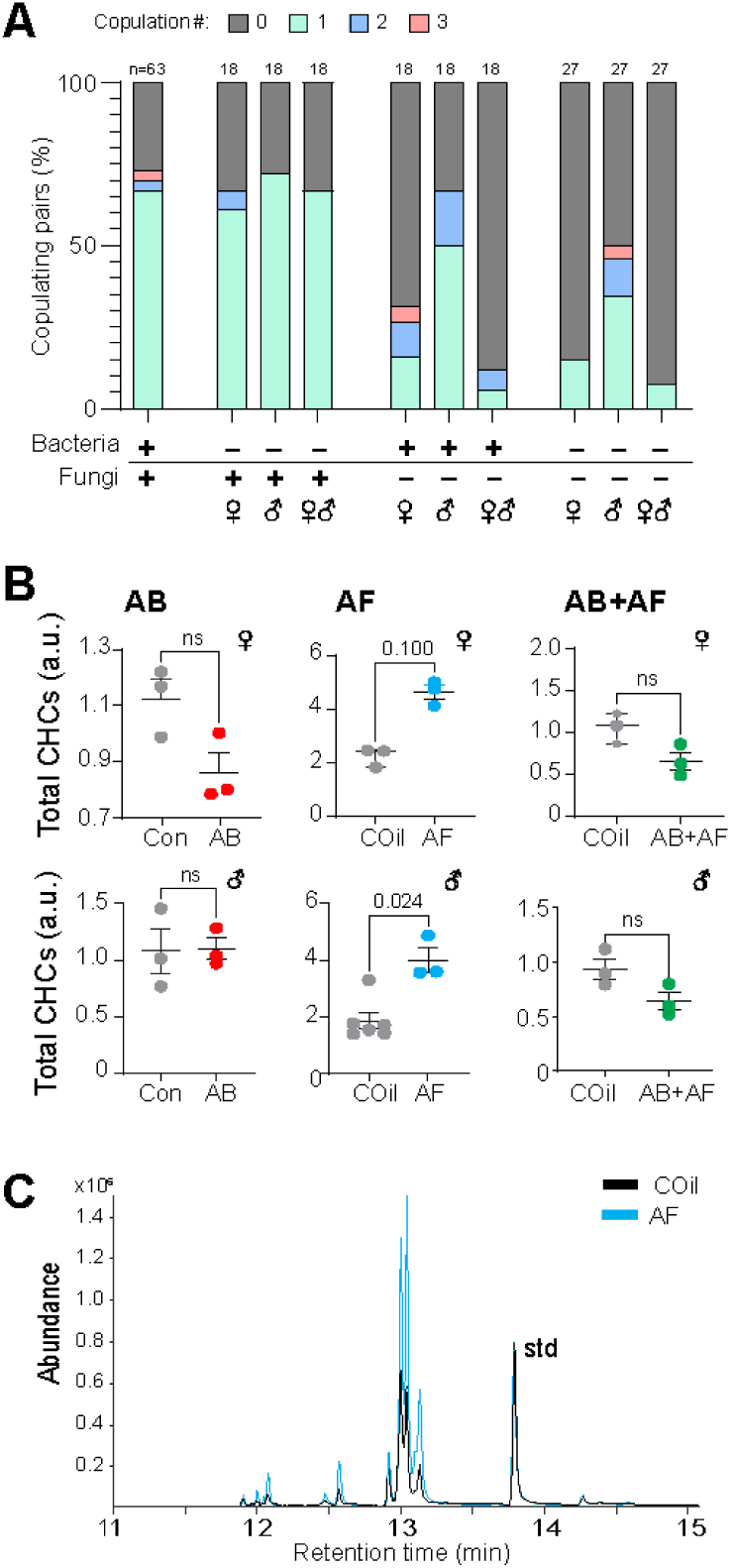
Impact of bacteria and fungi on mating and cuticular hydrocarbons. **A.** Proportion of females copulating 0, 1, 2, or 3 times during 48 hr mating trial. Copulation occurred with similar frequency amongst AB-treated flies. AF-treated females, but not males, were significantly less likely to mate (p=0.001). Mating amongst pairs with both AF-treated males and females was also significantly suppressed (p<.0001). Dyads involving AB+AF females copulated with lower frequency (p<0.0001) but AB+AF males had only a slight decrease in on mating frequency (p=0.049). Multiple copulations were more frequent in trials involving AF treated flies and in particular, AF males. Samples size is indicated above bars. The p-values were determined by a Fisher Exact Probability test. **B,** Cuticular hydrocarbon (CHC) total abundances. Overall levels of CHCs are higher in AF-treated females and males (n=3-6; Mann Whitney test). The CHC levels of AB and AB+AF-treated flies were unchanged by antimicrobial treatment; lines indicate mean ± S.E.M. Each replicate consists of extract from 3 flies. **C.** Representative GCMS chromatogram overlaying the CHC profile from an AF-treated female (blue) with a control (black) showing a notable difference in signal abundance for all major peaks. The chromatograms are normalized to a spiked standard (std).

In contrast to the limited effect of bacteria, the mycobiome had a considerably greater impact on female, but not male, mating drive. Significantly fewer females treated with antifungals (either alone or in combination with antibiotics) copulated compared with control females, regardless of the male treatment group (**Fig. 5A**). Additionally, a disproportionately low number of matings led to ovipositioning (AF: p=0.012, AB+AF: p<0.0001, Fisher Exact Probability test). By contrast, the frequency of male mating and of successful copulation did not change with captan treatment. In fact, fungal suppression in males either with AF or AB+AF treatments tended to increase instances of multiple copulations. Taken together, microbes impact the mating behaviors of females and males in distinct ways: female mating drive and successful copulation are influenced by both bacteria and fungi whereas male mating drive is largely unaffected by microbial contributions.

One important pre-copulatory trait that robustly influences the decision to court is pheromone signaling. For many insects, cuticular lipids function as sex pheromones that impel or inhibit the decision to mate. Interestingly, suppressing fungal communities results in a significant increase of both male and female cuticular hydrocarbon (CHC) levels (**Fig. 5B, C**). In other insect species, total CHC levels increase with age (Hugo et al., 2006; Ngumbi et al., 2020; Vernier et al., 2019). Indeed, *Drosophila melanogaster* use features of the CHC profile as an indicator of the age of the potential mate (Kuo et al., 2012). It remains to be tested whether the overall increase in CHC levels directly contributes to the reduction in mating drive observed in treated flies. Nonetheless, the observed difference in cuticular lipids indicates that in females, the microbiome has a systemic impact on multiple reproductive features of reproduction including ovary development and cuticular lipid levels, with the latter potentially serving as an honest indicator of fitness.

#### Fatty acid profiles

Reproduction is an expensive physiological process that requires a substantial investment of energy stores and can result in the diminution of life span for females (Chapman et al., 1995). As such, lipid metabolism and storage is intertwined with oogenesis and changes in lipid levels and composition are frequently associated with reproduction (Hansen et al., 2013). Given the robust effects that microbiome manipulation had on male and female fecundity, we next sought to determine whether whole body fatty acids (FA), a major storage source of energy, are also affected by the gut microbiome. For females, antibacterial treatments, which have a negligible impact on fecundity, had a slight but non-significant effect on fatty acid levels (**Fig. 6A**; **Supp. Table 1**). By contrast, AF and AB+AF treatments, both of which had sizeable effects on female reproduction, caused large changes in total FA levels, although in opposite directions (**Fig. 6A; Supp. Table 3 & 5**), and led to the shortening of fatty acid carbon chain length (**Supp. Fig. 3**). The most striking effects occurred with captan treatment which induced a near 2.5-fold increase in FAs. Interestingly, suppressing both bacteria and fungi, which eliminates almost all egg production and egg laying, resulted in a substantial decrease of FAs, a finding that contrasts with the outcomes of inhibiting either microbial kingdom alone. As with fecundity, interkingdom interactions and their metabolic products contribute to lipogenesis in a manner that is distinct from either kingdom alone.

**Figure 6.**
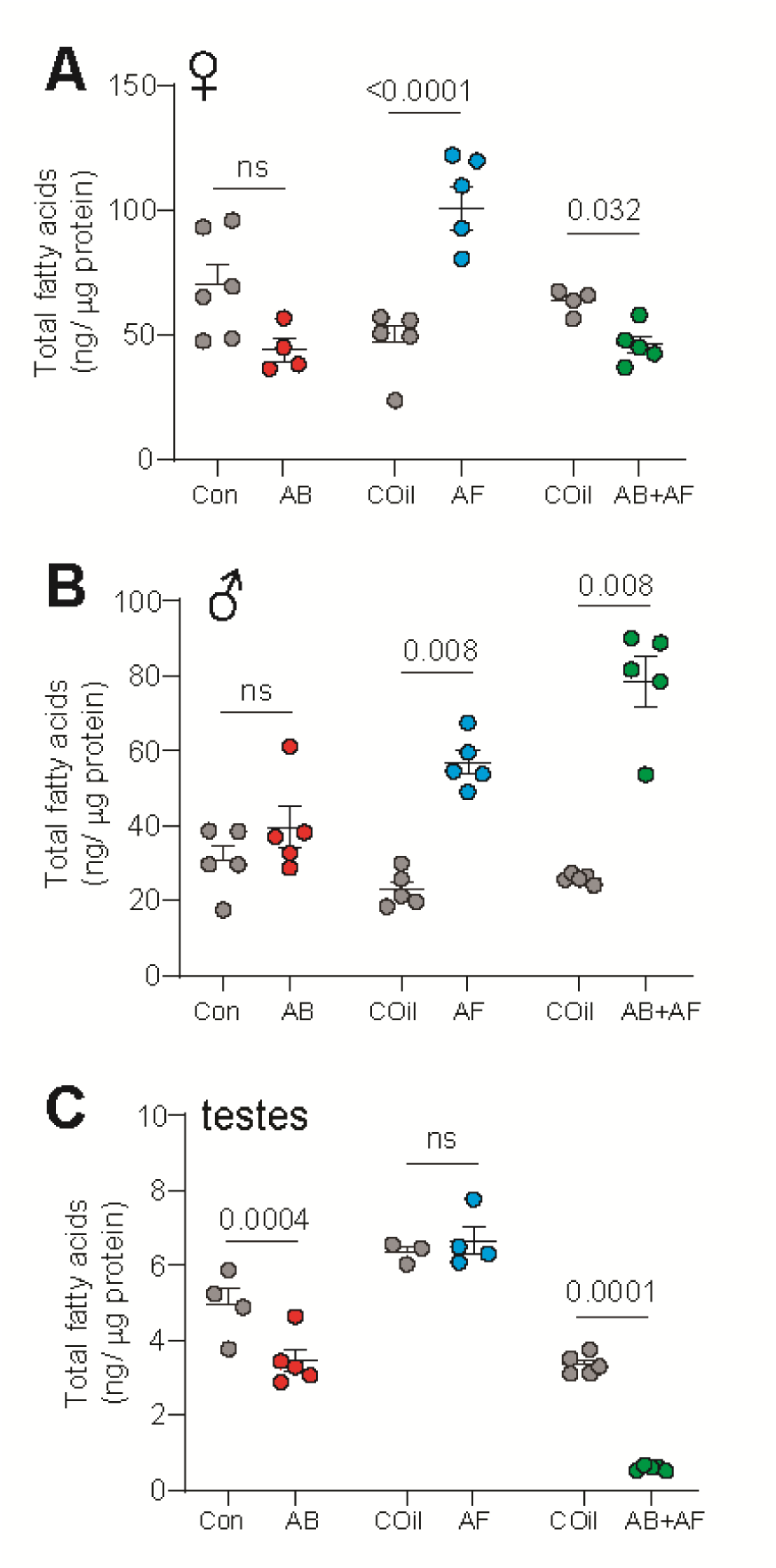
Impact of bacteria and fungi on the fatty acid content of the whole body and testes. **A.** Females treated with AB had similar levels of total fatty acid (FA) compared to controls (Mann Whitney test). AF-treated females contained significantly more total FAs compared to respective controls. AB+AF treatment resulted in a slight decree se of FAs; n=4-6 for all treatments; mean ± S.E.M. are shown. Each replicate consists of extract from 3 flies. **B.** Males treated with AB had similar total FA levels compared to controls (Mann Whitney test). Fatty acid content increased significantly with AF or AB+AF-treatment; n=5-6 for all treatments, mean ± S.E.M. are shown. Each replicate consists of extract from 3 flies. **C.** Testes of AB-treated males contained significantly lower FA levels (Mann-Whitney test). Unlike the results from whole body samples, FA levels in the testes were unchanged by AF treatment. The FA levels dropped significantly under AB+AF conditions; n=4-6 for all treatments, mean ± S.E.M. are shown. Each replicate consists of extract from 3 flies.

For males, no significant differences in whole body FA levels were found (**Supp. Table 2)** with AB treatment. The FA levels increased with captan administration, as seen with females (**Fig. 6B; Supp. Table 4**). In addition, the FA species shifted towards shorter carbon chain lengths for all treatments (**Supp. Fig, 4A-C**). However, in contrast to females, male FA levels increased substantially when both bacteria and fungi were suppressed (**Supp. Table 6**). We next asked whether the microbiome influences lipid metabolism in the testes. Fatty acids serve as useful indicators of reproductive health since they function as key components of phospholipids in the sperm cell membrane (Collodel et al., 2022; Gill & Valivety, 1997; Lenzi et al., 1996; Stubbs & Smith, 1984). In addition, Sertoli cells, a metabolic tissue in the testes, produces FA needed for germ cell maturation (Collodel et al., 2020; Esmaeili et al., 2015). Consistent with our findings that AB and AB+AF-treated flies exhibit reduced fecundity, FA levels in the testes decreased in response to both treatments (**Fig 6C**; **Supp. Table 7 & 9)**, possibly indicating a reduction in metabolic activity or lower levels of sperm production. In addition, FAs shortened in response to AB+AF treatment (**Supp. Fig 4F**) and unsaturation levels increased in response to AB treatment (**Supp. Fig 4J**). FA levels and profiles in the testes did not change with AF administration (**Fig. 6C; Supp. Fig 4 E, K**; **Supp. Table 8**). Overall, treatments that induced fecundity loss are associated with a reduction in testes FA levels. Moreover, the changes in testicular FA profiles are distinct from those observed in the whole body following antimicrobial treatment, indicating tissue-specific roles for the microbiome.

Changes in microbial community composition influence the composition and abundance of fatty acids throughout the body and reproductive tissues. However, females and males rely differently on fungi and bacteria for metabolic and reproductive needs. Although AB+AF treatment was detrimental for both male and female fecundity and mating (**Fig. 4 & 5**), the FA levels of males and females changed in opposite directions in the whole body when treated with AB+AF. This sex-specific contrast in lipid metabolic response is consistent with our observations that microbial interactions also exert sexually dimorphic effects on fecundity and mating drive.

## Discussion

Fungi and bacteria are prevalent in the microbiomes of many animals yet little is known about the contributions of each microbial kingdom or their interactions to host physiology (Belmonte et al., 2019; Douglas, 2018; McMullen et al., 2020; Murgier et al., 2019). By using antimicrobial drugs to separately and concurrently manipulate the populations of bacteria and fungi in an animal gut, we identified distinct contributions for each kingdom and their interactions in addition to discerning sexually dimorphic dependencies of the host on the microbiome for reproduction and lipid metabolism.

### Cross-kingdom co-dependency in the Drosophila microbiome

Axenic animals are standard experimental tools for elucidating the physiological function of the microbiome (Uzbay, 2019; Wu et al., 2023). For *Drosophila*, the process normally entails bleaching the eggs and maintaining a germ-free state throughout development. In contrast to generating axenic animals, a pharmaceutical-based strategy allows microbes to be conditionally suppressed only at the post-eclosion stage, thus avoiding developmental effects. The use of antimicrobial treatments also enabled the contributions of bacteria, fungi, and their interactions to be parsed. We note that the fungicide captan used in our study does not inhibit all fungal types and may also suppress some bacterial taxa (Banerjee & Banerjee, 1987; Rahden-Staroń et al., 1994). However, as indicated by CFU counts, each of the drug treatments effectively suppressed the targeted microbial populations and shifted the microbiome profile in terms of taxonomic diversity and bacterial composition. Interestingly, suppressing fungal growth also led to a change in the community profile of bacteria (**Fig. 2**). *Enterococcus* populations expanded when fungal populations are suppressed, indicating that in a balanced microbiome, fungi inhibit overgrowth of these taxa. This finding is consistent with a previous study in mammals showing that the administration of antifungal drugs in a model of mouse colitis resulted in an expansion of bacterial genera (Qiu et al., 2015). We hypothesize that bacterial-fungal interactions help to maintain community structure, possibly by providing physical scaffolding, availability of critical cofactors, or antagonizing the proliferation of particular bacteria (Biedermann & Vega, 2020; Li et al., 2021; Pierce et al., 2021).

In contrast to the bacterial community, fungal composition was substantially more resilient. No significant changes in beta diversity were detected under any of the treatment conditions although *Saccharomycopsis* fungi increased with antibacterial treatments. Overall, our manipulations of the *Drosophila* bacterial-fungal microbial community reveal that whilst bacteria composition depends on the fungal community, fungi composition is relatively stable despite changes in bacteria. This finding runs contrary to the human gut mycobiome where, for the most part, fungi are thought to be more transitory and variable than the bacterial microbiome (Hallen-Adams & Suhr, 2017; Nash et al., 2017; Santus et al., 2021). However, some studies indicate a stable, resident mycobiome co-exists with the transient mycobiome (Fiers et al., 2019; Monteiro-da-Silva et al., 2014; Nash et al., 2017; Shuai et al., 2022; Szóstak et al., 2023). Determining whether a stable mycobiome is a feature specific to insects that harbor commensal fungi or a general rule of organismal microbiomes will require more bacterial/ fungal manipulation studies across other taxonomic levels.

### Sexually dimorphic roles of the microbiome in fecundity

Previous studies of *Drosophila* have established that gut bacteria play a significant role in life history features (Douglas, 2019; Lee et al., 2019; Matthews et al., 2021). Notably, changes in the taxonomic diversity of the microbiome as well as the interplay between bacteria influence the tradeoff between lifespan and reproduction in females (Gould et al., 2018; Shu et al., 2021; Walters et al., 2020). Our results reveal that interactions between bacteria and fungi are similarly important for both male and female reproduction since suppressing both kingdoms resulted in a greater loss of fecundity than suppressing either one alone. Additionally, bacteria and fungi alter male and female reproduction in different ways. Bacteria is needed by both sexes for successful copulation, as measured by oviposition frequency following mating (**Fig. 4E**). Male mating success is also affected by the loss of bacteria but not fungal communities. By contrast, fungi are more critical for female fecundity, contributing to oogenesis, mating drive, copulation success, and oviposition behavior. Each of these features were significantly diminished with AF treatment. Metabolically active cells appear to be necessary for oogenesis enhancement: only active fecal transfers were capable of partially rescuing female fecundity (**Fig. 3D**). Moreover, fecundity could be rescued by co-housing AB+AF-treated flies with control flies but not AB+AF-treated flies. This outcome highlights the importance of microbiome composition rather than biomass in reversing the deleterious effects of microbiome suppression on female oogenesis.

The sexual dimorphism in host-microbe interactions may partly be explained by the different responses of each sex to antifungal treatment. In particular, *Acetobacter* and *Lactobacillus*, both of which have been linked to the fertility of *D. melanogaster* ((Elgart et al., 2016; Pais et al., 2018)Gould et al. 2018), were substantially reduced in females compared to males. However, antibacterial treatment, which also reduces the relative abundances of *Acetobacter* and *Lactobacillus (***Fig. 2**), had only slight effects on female fecundity and oogenesis, indicating that captan-induced changes in the bacterial profile is not the primary cause of fecundity. Interestingly, Gould et al. (2018) were not able to rescue fecundity in germ-free female *D. melanogaster* after providing these flies with a mix of healthy bacteria, alluding to the possibility that fungi are needed to fully rescue female *Drosophila* fecundity. Taken together, these outcomes reveal direct roles for fungal activity in female reproduction in addition to maintaining bacterial community stability, ensuring the presence of key beneficial microbes, and providing nutrition (Drummond-Barbosa & Spradling, 2001).

The reduction in female mating may either be due to an increased reluctance to mate or a decreased attractiveness to males. Female copulation can incur a fitness cost including infection from microbes transferred in male ejaculate, genital damage and early death (Crudgington & Siva-Jothy, 2000; Rowe et al., 2020; Short et al., 2012). As such, limiting mating activity in the absence of ovary development or a compromised microbial defense system may be a reproductive strategy to conserve resources and minimize physical harm. Although female CHCs levels increased with captan treatment, we did not test whether males perceive and respond differently to the change. A more detailed analysis of courtship behavior will be needed to assess whether female sex appeal has changed with AF treatment.

### How do microbes influence fecundity?

Lipid metabolism and reproduction are mechanistically coupled through numerous shared metabolic underpinnings (Hansen et al., 2013). Our studies show that the microbiome is a central modulator of these systemic traits: alterations in microbe composition are associated with changes in fecundity, fatty acid levels, and fatty acid composition in the whole body as well as the testes. Moreover, bacteria, fungi, and bacterial-fungal interactions alter the lipid profiles in distinct ways, indicating that the changes in FAs are not simply due to a loss of nutrition. Previous studies have shown that microbes supply FA precursors and that lipid profiles change when bacteria are eliminated (Schwenke et al., 2016). Alterations in female FA levels likely reflect a homeostatic shift between fat storage and reproduction, potentially mediated by circulating hormones such as ecdysone, insulin, or juvenile hormone (Nässel & Zandawala, 2020). As for male fecundity, we observed that FA levels in the testes dropped significantly with the suppression of bacteria or both bacteria and fungi. Precursors supplied by bacteria may be incorporated into the biosynthesis of sperm membranes. Alternatively, energy stores could be shunted from spermatogenesis or accessory gland protein production to support other physiological needs such as immune defense (Fedorka et al., 2004; Gupta et al., 2013; McKean & Nunney, 2001; Schwenke et al., 2016), telomere length (indicator of aging and lifespan) (Morbiato et al., 2023) and defensive weapons (Cavender et al., 2021).

In addition to changing patterns of lipid storage, microbes also influence the profile of FA species used by the host. Shifts in FA length can alter functional properties of spermatozoa including membrane fluidity and sperm motility (Stubbs & Smith, 1984). Here we show that microbes alter lipid metabolism in a tissue-specific manner resulting in different lipid profiles in the testes compared to the whole body and these alterations are associated with a loss of male fecundity.

### Implications for the rapid radiation of Hawaiian Drosophila

*Drosophila*, like many animals, rely on external sources such as diet to renew their microbiome (Blum et al., 2013; Chandler et al., 2011; Staubach et al., 2013). The diversity of yeast associate with Hawaiian PWDs and their host plants has been proposed as a factor in PWD rapid speciation. Given our findings that the mycobiome is coupled to female fecundity and mating drive, variation in fungal communities either through host plant selection or abiotic conditions may contribute to reproductive isolation. In addition to fecundity, our results reveal that CHC levels are also influenced by microbe composition. Considering that CHCs serve as a barrier to desiccation, these outcomes have fascinating implications for the role of microbes in enabling host tolerance to local temperature and humidity conditions. As such, microbes acquired from host-plants may have contributed to PWDs explosive radiation across the Hawaiian archipelago by facilitating rapid adaptation. Future experiments measuring the interaction between microbe composition and temperature and humidity tolerance under natural conditions are needed to address this prediction.

## Conclusion

Our results reveal distinct and sexually dimorphic roles for the microbiome with respect to fecundity, mating behavior, and lipid metabolism. Fungal metabolic activity supports female oogenesis, influences mating drive, and helps to maintain the presence of key beneficial bacterial taxa. By contrast, gut bacteria support male fecundity whilst fungi have limited contributions. For both sexes, reproductive fitness depends critically on a balance between bacterial and fungal components of the microbiome and their interactions. Upsetting this equilibrium can have devastating consequences on male and female reproduction.

## Acknowledgements

This work was funded by the National Science Foundation Grant No. 2025669 (JYY), National Institute of General Medical Sciences of the National Institutes of Health Grant No. P20GM125508 (JYY), P20GM103466 (JYY), and Hawaiʻi Community Foundation Grant No. 19CON-95452 (JYY). We are grateful to Margaret McFall-Ngai, Edward Ruby, Sean Schoville, Sean Swift, Mélisandre Téfit, Laura Tipton, Nicolas Cetraro, Kelli Konicek for helpful scientific discussions. We thank Leon Boyles and David Zheng for excellent technical assistance. All GCMS measurements and HTS library preparations were performed in the UHM Microbial Genomics and Analytical Laboratory core (supported by P20GM125508). Sequencing was performed at the UHM Advanced Studies in Genomics, Proteomics and Biooinformatics facility. Microscopy images were acquired in the UHM Microscopy Imaging Center for Research through Observation (supported by P20GM139753).

## Author Contributions

MJM designed and performed research, analyzed data, and contributed to the writing and editing of the manuscript. LS and AM performed research and analyzed data. DKP provided resources, organized the raising of *D. grimshawi* used in this study, and contributed to editing of the paper. JYY designed and performed research, contributed resources, analyzed data, and contributed to the writing and editing of the manuscript.

---

**Supplemental Figure 1.**
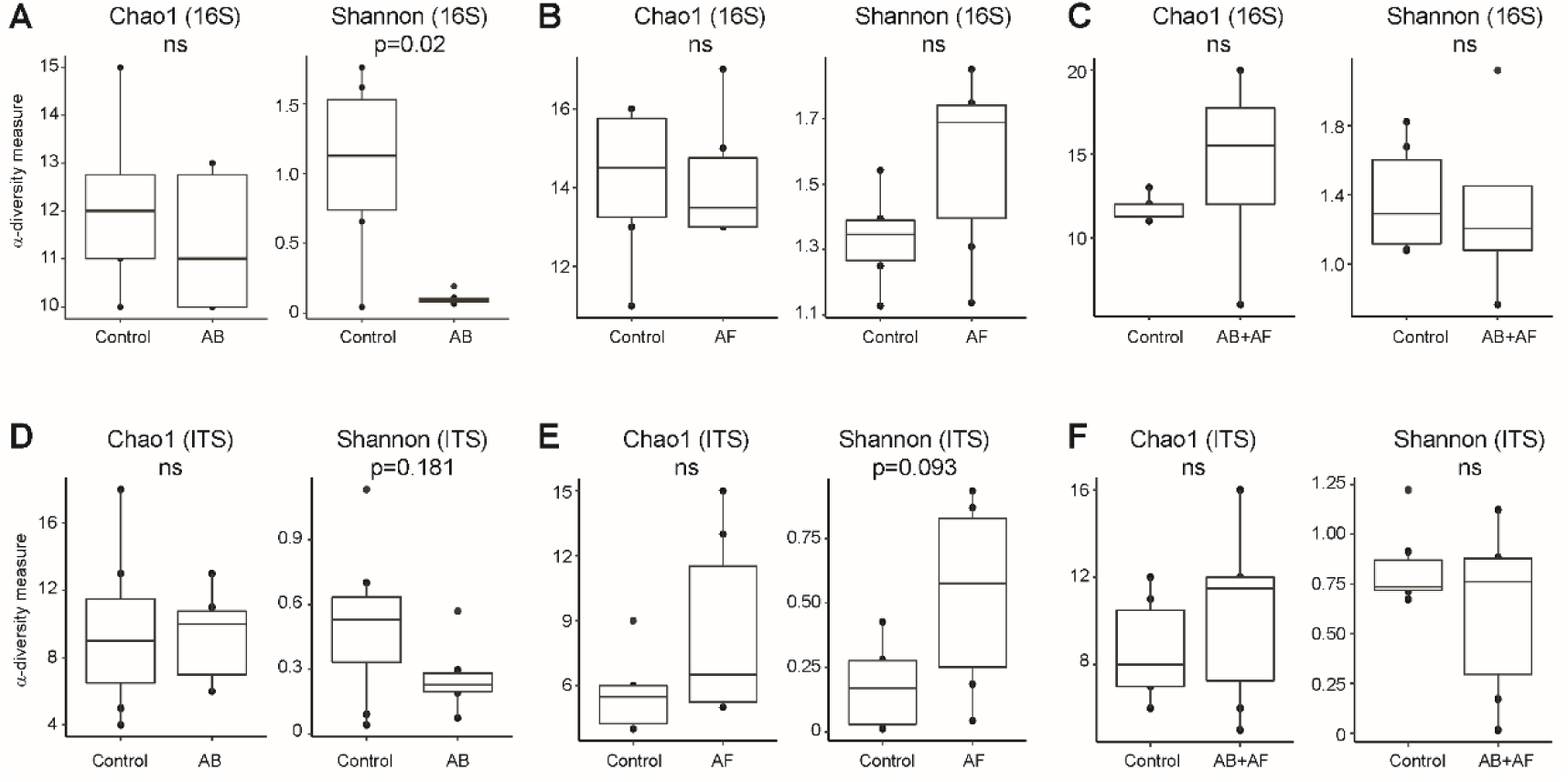
**A – F**. Chao1 and Shannon α-diversity indices of bacterial (top row) and fungal communities (bottom row) in control and antimicrobial-treated flies. α-diversity was compared using a Wilcoxon rank-sum test, n=6-8 for all treatments; ns: not significant (p>0.05).

**Supplemental Figure 2.**
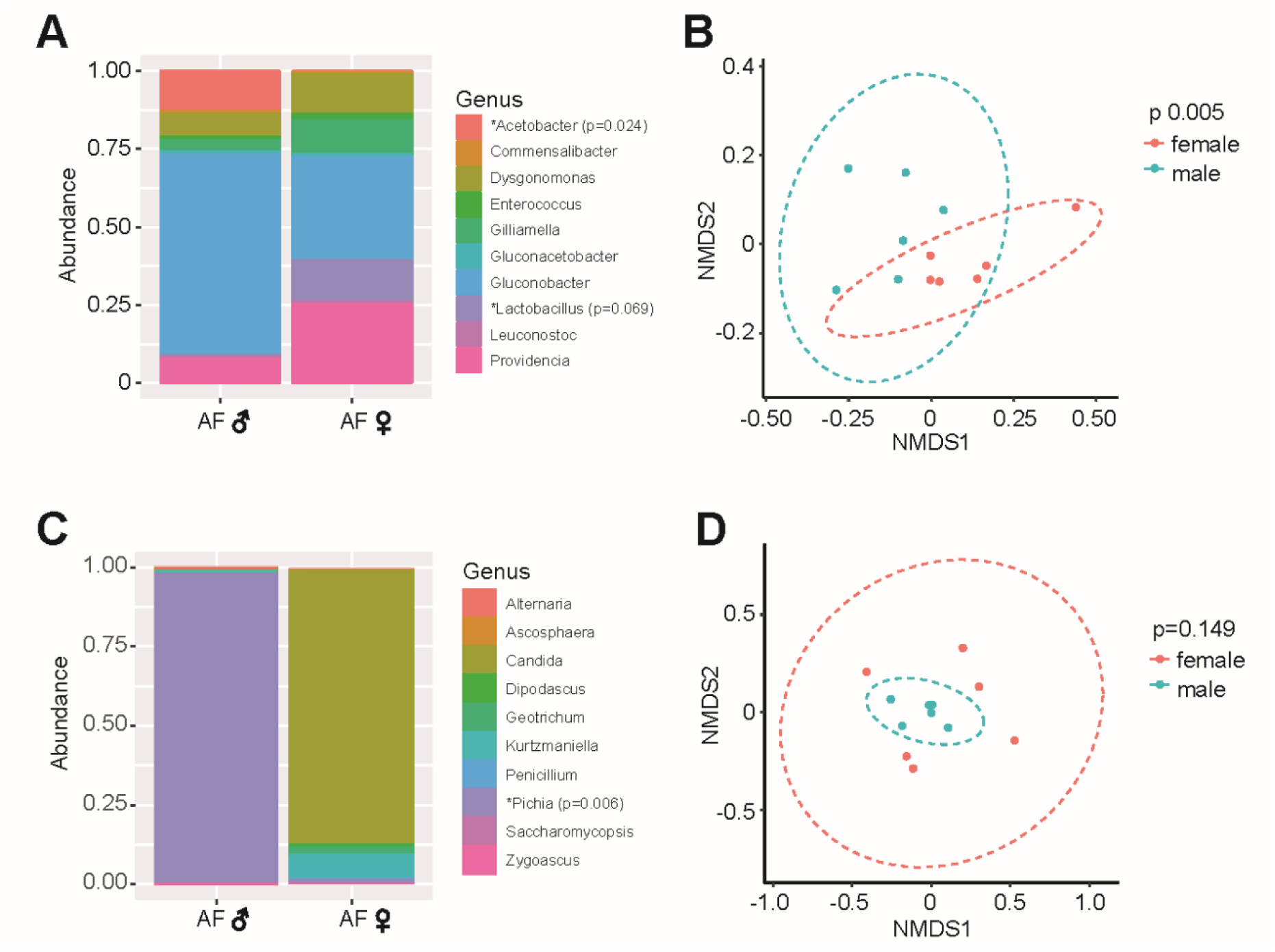
Comparison of male and female microbiome 16S and ITS high throughput sequencing amplicon profiles following antifungal treatment. Samples are comprised of flies that were treated for 21 and 35 days. **A.** Scaled relative abundance plots for the 10 most abundant bacterial genera of males and females following antifungal (AF) treatment (n=6 per sex). The p values were determined using univariate multiple testing with an F test; *: abundance is significantly different between control and treatment conditions. **B.** Non-multidimensional scaling plots (NMDS; based on Jaccard distances) of OTUs reveal distinct bacterial communities in males and females following AF treatment (ANOSIM, n=6 per sex). Ellipses represent significance at 0.05 confidence. **C.** Scaled relative abundance plots for the 10 most abundant fungal genera of males and females following antifungal (AF) treatment (n=6 per sex). The p values were determined using univariate multiple testing with an F test; *: significantly different between control and treatment conditions. **D.** Non-multidimensional scaling plots (NMDS; based on Jaccard distances) of OTUs show no significant separation in fungal composition between males and females (ANOSIM, n=6 per sex). Ellipses represent significance at 0.05 confidence.

**Supplemental Figure 3.**
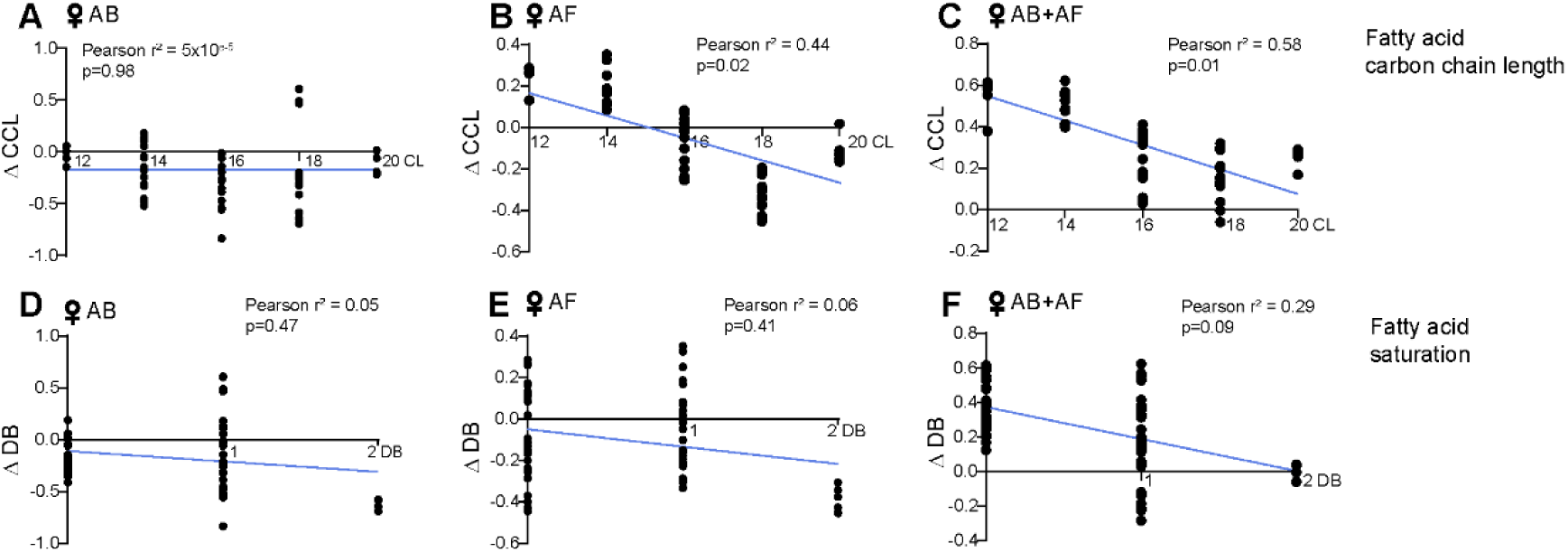
Influence of microbiome composition on female whole body fatty acid carbon chain length and degree of saturation. The Pearson r^2^ value and simple linear regression line are provided; n=3-5. **A – C.** Change in whole body fatty acid carbon chain lengths (CCL) in antibacterial-(AB), antifungal-(AF), or AB+AF treated-females compared to controls. **D – F.** Change in whole body fatty acid saturation levels in treated females compared to controls; DB: double bond number.

**Supplemental Figure 4.**
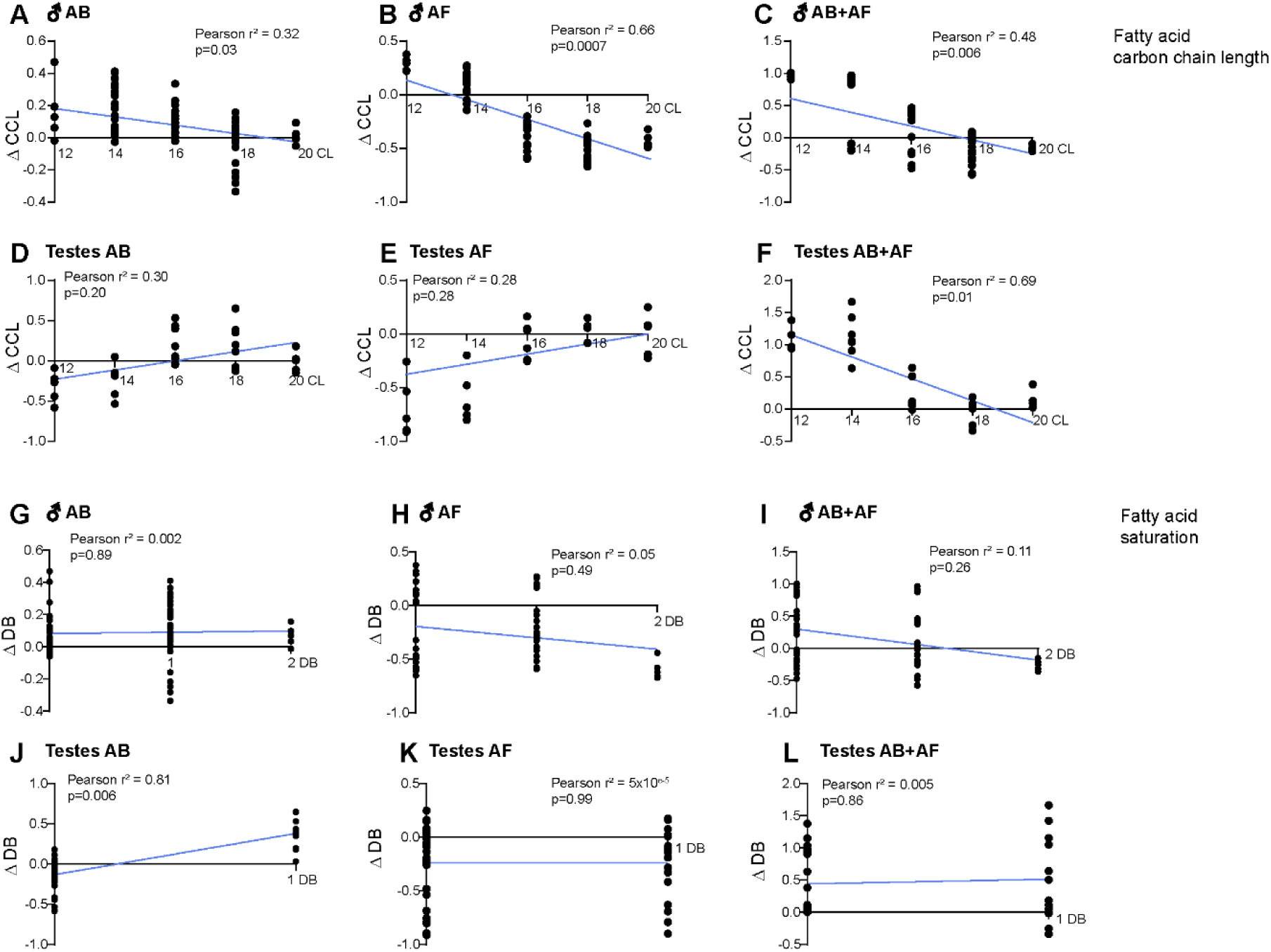
Influence of microbiome composition on male whole body and testes fatty acid carbon chain length and degree of saturation. The Pearson r^2^ value and simple linear regression line are provided; n=3-5. **A – C.** Change in whole body fatty acid carbon chain lengths (CCL) in antibacterial (AB), antifungal (AF), or AB+AF treated-males compared to controls. **D – F.** Change in testes fatty acid carbon chain lengths (CCL) in antibacterial-(AB), antifungal-(AF), or AB+AF-treated males compared to controls. **G – I.** Change in whole body fatty acid saturation levels in treated males compared to controls; DB: double bond number. **J – L.** Change in testes fatty acid saturation levels in treated males compared to controls; DB: double bond number.

## Notes

### Competing Interest Statement

The authors have declared no competing interest.

### Summary of Updates

Citations corrected.

