## Supplemental Tables for "Bacterial and fungal components of the gut microbiome have distinct, sex-specific roles in Hawaiian *Drosophila* reproduction"

**Supplemental Table 1.** Fatty acid (FA) content of control and antibacterial (AB)-treated females for each replicate.

| | Control[3] | | | | | AB[3] | | | | |
|---|---|---|---|---|---|---|---|---|---|---|
| **FA[1,2]** | **1** | **2** | **3** | **4** | **5** | **1** | **2** | **3** | **4** | **5** |
| C12:0 | 7.065 | 14.867 | 6.982 | 16.838 | 11.414 | 10.490 | 4.955 | 4.065 | 4.541 | 8.127 |
| C13:0 | 0.000 | 0.007 | 0.000 | 0.008 | 0.006 | 0.021 | 0.005 | 0.013 | 0.007 | 0.019 |
| C14:2 | 0.000 | 0.000 | 0.000 | 0.014 | 0.000 | 0.048 | 0.034 | 0.030 | 0.034 | 0.035 |
| C14:1a | 0.228 | 0.418 | 0.261 | 0.577 | 0.519 | 0.718 | 0.454 | 0.419 | 0.330 | 0.584 |
| C14:1b | 0.298 | 0.630 | 0.358 | 0.541 | 0.414 | 0.365 | 0.199 | 0.147 | 0.183 | 0.262 |
| C14:1c | 1.727 | 3.693 | 1.842 | 3.700 | 2.557 | 1.847 | 0.934 | 0.537 | 0.791 | 1.334 |
| C14:0 | 8.727 | 19.662 | 7.362 | 21.489 | 15.970 | 15.950 | 7.959 | 7.231 | 6.827 | 13.608 |
| C16:1a | 0.050 | 0.091 | 0.026 | 0.075 | 0.085 | 0.133 | 0.085 | 0.092 | 0.130 | 0.120 |
| C16:1b | 2.957 | 5.606 | 2.624 | 5.264 | 4.116 | 3.302 | 1.933 | 1.560 | 2.068 | 2.773 |
| C16:0 | 6.246 | 12.740 | 6.149 | 13.205 | 10.163 | 10.955 | 7.045 | 6.994 | 8.022 | 10.142 |
| C17:0 | 0.028 | 0.021 | 0.017 | 0.034 | 0.021 | 0.028 | 0.020 | 0.030 | 0.020 | 0.030 |
| C18:2 | 0.104 | 0.173 | 0.109 | 0.191 | 0.126 | 0.293 | 0.285 | 0.336 | 0.279 | 0.294 |
| C18:1a | 2.156 | 3.546 | 2.334 | 3.452 | 2.813 | 2.957 | 2.300 | 2.283 | 2.726 | 2.797 |
| C18:1b | 0.142 | 0.217 | 0.170 | 0.197 | 0.148 | 0.074 | 0.040 | 0.010 | 0.005 | 0.077 |
| C18:0 | 0.914 | 1.192 | 1.009 | 1.309 | 1.099 | 1.168 | 0.987 | 0.916 | 0.985 | 1.389 |
| C20:0 | 0.148 | 0.194 | 0.150 | 0.160 | 0.121 | 0.118 | 0.091 | 0.103 | 0.092 | 0.130 |
| C22:0 | 0.021 | 0.019 | 0.010 | 0.018 | 0.013 | 0.015 | 0.009 | 0.011 | 0.005 | 0.015 |

[1]Notation indicates carbon chain length followed by number of double bonds. Positional isomers for C14:1, C16:1, and C18:1 FA species are indicated by “a”, b”, and “c”.
[2]Each replicate is comprised of homogenate from 5 flies.
[3]Raw abundance counts are normalized to the abundance of a spiked standard (pentadecanoic acid).

**Supplemental Table 2.** Fatty acid (FA) content of control and antibacterial (AB)-treated males for each replicate.

| | Control[3] | | | | | AB[3] | | | | |
|---|---|---|---|---|---|---|---|---|---|---|
| **FA[1,2]** | **1** | **2** | **3** | **4** | **5** | **1** | **2** | **3** | **4** | **5** |
| C10:0 | 0.023 | 0.020 | 0.014 | 0.000 | 0.016 | 0.037 | 0.026 | 0.013 | 0.020 | 0.023 |
| C12:0 | 3.983 | 3.124 | 2.684 | 0.820 | 2.894 | 8.086 | 3.626 | 1.999 | 3.463 | 3.393 |
| C13:0 | 0.022 | 0.012 | 0.018 | 0.000 | 0.013 | 0.032 | 0.015 | 0.000 | 0.013 | 0.027 |
| C14:1a | 0.549 | 0.211 | 0.368 | 0.193 | 0.340 | 0.821 | 0.567 | 0.276 | 0.393 | 0.577 |
| C14:1b | 0.239 | 0.183 | 0.158 | 0.081 | 0.162 | 0.363 | 0.338 | 0.247 | 0.310 | 0.365 |
| C14:1c | 0.756 | 0.656 | 0.525 | 0.165 | 0.626 | 1.429 | 0.588 | 0.497 | 0.688 | 0.675 |
| C14:0 | 7.218 | 6.113 | 4.805 | 1.863 | 5.638 | 13.563 | 6.872 | 3.829 | 6.647 | 6.263 |
| C16:1a | 0.139 | 0.084 | 0.109 | 0.075 | 0.108 | 0.248 | 0.181 | 0.089 | 0.115 | 0.161 |
| C16:1b | 2.170 | 2.101 | 1.815 | 1.105 | 1.874 | 3.424 | 2.355 | 2.106 | 2.613 | 2.317 |
| C16:0 | 9.896 | 10.816 | 7.957 | 4.916 | 8.358 | 13.568 | 9.800 | 6.714 | 8.666 | 9.528 |
| C17:0 | 0.065 | 0.050 | 0.052 | 0.041 | 0.052 | 0.059 | 0.058 | 0.045 | 0.048 | 0.063 |
| C18:2 | 0.269 | 0.254 | 0.236 | 0.181 | 0.209 | 0.373 | 0.301 | 0.249 | 0.205 | 0.303 |
| C18:1a | 3.071 | 2.617 | 2.579 | 1.936 | 2.447 | 3.774 | 2.838 | 2.415 | 2.539 | 2.883 |
| C18:1b | 0.187 | 0.177 | 0.131 | 0.079 | 0.124 | 0.106 | 0.079 | 0.066 | 0.075 | 0.076 |
| C18:0 | 1.428 | 1.411 | 1.228 | 1.063 | 1.242 | 1.588 | 1.368 | 1.268 | 1.213 | 1.357 |
| C20:0 | 0.241 | 0.232 | 0.241 | 0.176 | 0.223 | 0.266 | 0.221 | 0.239 | 0.200 | 0.269 |
| C22:0 | 0.044 | 0.049 | 0.069 | 0.060 | 0.054 | 0.053 | 0.050 | 0.065 | 0.052 | 0.077 |

[1]Notation indicates carbon chain length followed by number of double bonds. Positional isomers for C14:1, C16:1, and C18:1 FA species are indicated by “a”, b”, and “c”.
[2]Each replicate is comprised of homogenate from 5 flies.
[3]Raw abundance counts are normalized to the abundance of a spiked standard (pentadecanoic acid).

**Supplemental Table 3.** Fatty acid (FA) content of control and antifungal (AF)-treated females for each replicate.

| | Control[3] | | | | | AF[3] | | | | |
|---|---|---|---|---|---|---|---|---|---|---|
| **FA[1,2]** | **1** | **2** | **3** | **4** | **5** | **1** | **2** | **3** | **4** | **5** |
| C10:0 | 0.104 | 0.150 | 0.000 | 0.189 | 0.000 | 0.000 | 0.104 | 0.000 | 0.099 | 0.041 |
| C12:0 | 12.266 | 22.722 | 9.747 | 16.916 | 8.545 | 12.991 | 15.785 | 10.605 | 17.472 | 10.048 |
| C13:0 | 0.023 | 0.039 | 0.015 | 0.022 | 0.013 | 0.030 | 0.030 | 0.017 | 0.033 | 0.019 |
| C14:2 | 0.031 | 0.046 | 0.040 | 0.028 | 0.024 | 0.042 | 0.026 | 0.000 | 0.042 | 0.022 |
| C14:1a | 0.550 | 0.923 | 0.720 | 0.699 | 0.466 | 0.263 | 0.288 | 0.190 | 0.341 | 0.198 |
| C14:1b | 0.369 | 0.612 | 0.438 | 0.423 | 0.337 | 0.337 | 0.360 | 0.257 | 0.397 | 0.226 |
| C14:1c | 2.067 | 0.000 | 1.497 | 2.646 | 1.193 | 3.183 | 3.767 | 2.535 | 3.501 | 2.156 |
| C16:1a | 0.107 | 0.125 | 0.114 | 0.122 | 0.078 | 0.091 | 0.071 | 0.072 | 0.087 | 0.054 |
| C16:1b | 3.147 | 5.166 | 2.592 | 4.194 | 2.181 | 4.079 | 4.706 | 3.115 | 4.793 | 2.385 |
| C16:0 | 9.728 | 14.476 | 8.264 | 12.099 | 7.062 | 9.990 | 10.940 | 7.174 | 11.202 | 5.828 |
| C17:0 | 0.050 | 0.065 | 0.049 | 0.050 | 0.036 | 0.030 | 0.028 | 0.018 | 0.027 | 0.022 |
| C18:2 | 0.174 | 0.251 | 0.185 | 0.217 | 0.150 | 0.140 | 0.117 | 0.105 | 0.128 | 0.092 |
| C18:1a | 2.401 | 3.353 | 2.549 | 2.990 | 1.989 | 2.160 | 2.278 | 1.811 | 2.505 | 1.368 |
| C18:1b | 0.092 | 0.138 | 0.086 | 0.107 | 0.082 | 0.080 | 0.075 | 0.000 | 0.073 | 0.051 |
| C18:0 | 1.248 | 1.581 | 1.409 | 1.364 | 0.963 | 0.890 | 0.935 | 0.811 | 0.965 | 0.741 |
| C19:0 | 0.028 | 0.042 | 0.036 | 0.026 | 0.017 | 0.025 | 0.022 | 0.000 | 0.021 | 0.014 |
| C20:0 | 0.403 | 0.477 | 0.364 | 0.439 | 0.285 | 0.359 | 0.308 | 0.229 | 0.332 | 0.234 |
| C22:0 | 0.143 | 0.140 | 0.109 | 0.151 | 0.097 | 0.119 | 0.101 | 0.082 | 0.101 | 0.083 |

[1]Notation indicates carbon chain length followed by number of double bonds. Positional isomers for C14:1, C16:1, and C18:1 FA species are indicated by “a”, b”, and “c”.
[2]Each replicate is comprised of homogenate from 5 flies.
[3]Raw abundance counts are normalized to the abundance of a spiked standard (pentadecanoic acid).

**Supplemental Table 4.** Fatty acid (FA) content of control and antibacterial (AB)-treated males for each replicate.

| | Control[3] | | | | | AF[3] | | | | |
|---|---|---|---|---|---|---|---|---|---|---|
| **FA[1,2]** | **1** | **2** | **3** | **4** | **5** | **1** | **2** | **3** | **4** | **5** |
| C10:0 | 0.000 | 0.034 | 0.000 | 0.000 | 0.040 | 0.128 | 0.084 | 0.126 | 0.128 | 0.099 |
| C12:0 | 1.237 | 2.106 | 1.538 | 2.049 | 2.912 | 18.338 | 11.392 | 15.394 | 14.043 | 12.401 |
| C13:0 | 0.000 | 0.000 | 0.000 | 0.000 | 0.035 | 0.027 | 0.000 | 0.000 | 0.000 | 0.021 |
| C14:1a | 0.163 | 0.261 | 0.055 | 0.086 | 0.098 | 0.436 | 0.300 | 0.340 | 0.319 | 0.323 |
| C14:1b | 0.105 | 0.121 | 0.061 | 0.093 | 0.094 | 0.430 | 0.336 | 0.383 | 0.289 | 0.361 |
| C14:1c | 0.219 | 0.222 | 0.235 | 0.320 | 0.496 | 2.371 | 1.590 | 1.879 | 1.298 | 2.064 |
| C14:0 | 1.647 | 2.906 | 2.126 | 2.893 | 3.807 | 15.036 | 9.881 | 11.775 | 11.224 | 13.129 |
| C16:1a | 0.061 | 0.073 | 0.039 | 0.048 | 0.062 | 0.126 | 0.097 | 0.094 | 0.109 | 0.115 |
| C16:1b | 1.015 | 1.205 | 1.026 | 1.302 | 1.631 | 2.910 | 2.229 | 2.375 | 1.908 | 2.673 |
| C16:0 | 4.831 | 6.299 | 4.662 | 5.684 | 7.136 | 7.786 | 5.180 | 6.122 | 6.089 | 7.336 |
| C17:0 | 0.040 | 0.049 | 0.032 | 0.030 | 0.046 | 0.028 | 0.000 | 0.026 | 0.025 | 0.030 |
| C18:2 | 0.203 | 0.269 | 0.131 | 0.140 | 0.213 | 0.205 | 0.143 | 0.167 | 0.209 | 0.190 |
| C18:1a | 1.736 | 2.361 | 1.343 | 1.586 | 1.817 | 2.115 | 1.675 | 1.843 | 1.839 | 2.064 |
| C18:1b | 0.105 | 0.152 | 0.083 | 0.095 | 0.114 | 0.152 | 0.158 | 0.184 | 0.181 | 0.159 |
| C18:0 | 0.874 | 1.083 | 0.859 | 0.861 | 0.983 | 0.966 | 0.870 | 0.947 | 0.998 | 0.970 |
| C20:0 | 0.196 | 0.233 | 0.136 | 0.162 | 0.173 | 0.266 | 0.222 | 0.249 | 0.263 | 0.274 |
| C22:0 | 0.064 | 0.072 | 0.045 | 0.051 | 0.069 | 0.069 | 0.066 | 0.072 | 0.070 | 0.072 |

[1]Notation indicates carbon chain length followed by number of double bonds. Positional isomers for C14:1, C16:1, and C18:1 FA species are indicated by “a”, b”, and “c”.
[2]Each replicate is comprised of homogenate from 5 flies.
[3]Raw abundance counts are normalized to the abundance of a spiked standard (pentadecanoic acid).

**Supplemental Table 5.** Fatty acid (FA) content of control and antibacterial and antifungal (AB+AF)-treated females for each replicate.

| | Control[3] | | | | | AB+AF[3] | | | | |
|---|---|---|---|---|---|---|---|---|---|---|
| **FA[1,2]** | **1** | **2** | **3** | **4** | **5** | **1** | **2** | **3** | **4** | **5** |
| C12:0 | 8.412 | 17.665 | 17.424 | 15.381 | 17.827 | 18.195 | 11.926 | 15.477 | 15.780 | 13.357 |
| C14:1a | 2.308 | 1.388 | 1.107 | 0.953 | 1.102 | 0.288 | 0.170 | 0.236 | 0.268 | 0.222 |
| C14:1b | 1.843 | 0.690 | 0.645 | 0.497 | 0.553 | 0.271 | 0.166 | 0.210 | 0.204 | 0.177 |
| C14:1c | 6.579 | 2.969 | 2.905 | 2.047 | 2.486 | 3.800 | 2.304 | 3.038 | 2.677 | 2.896 |
| C14:0 | 10.557 | 18.048 | 18.498 | 16.274 | 18.385 | 16.430 | 11.245 | 12.892 | 12.243 | 11.991 |
| C16:1a | 0.588 | 0.217 | 0.206 | 0.198 | 0.179 | 0.133 | 0.061 | 0.100 | 0.071 | 0.081 |
| C16:1b | 14.969 | 4.492 | 4.498 | 3.606 | 3.988 | 4.239 | 2.394 | 3.216 | 2.844 | 2.890 |
| C16:0 | 11.079 | 14.910 | 14.011 | 11.904 | 13.210 | 10.564 | 5.994 | 8.865 | 7.476 | 7.969 |
| C17:0 | 0.000 | 0.063 | 0.052 | 0.050 | 0.059 | 0.000 | 0.000 | 0.000 | 0.000 | 0.000 |
| C18:2 | 7.345 | 0.346 | 0.307 | 0.294 | 0.310 | 0.200 | 0.135 | 0.170 | 0.150 | 0.145 |
| C18:1a | 26.397 | 3.832 | 3.717 | 3.034 | 3.373 | 2.574 | 1.461 | 2.244 | 1.966 | 1.714 |
| C18:1b | 1.236 | 0.178 | 0.175 | 0.155 | 0.126 | 0.073 | 0.000 | 0.062 | 0.062 | 0.000 |
| C18:0 | 2.344 | 2.391 | 2.206 | 1.711 | 1.891 | 1.077 | 0.987 | 1.222 | 0.990 | 0.840 |
| C19:0 | 0.000 | 0.000 | 0.000 | 0.000 | 0.000 | 0.000 | 0.168 | 0.000 | 0.000 | 0.000 |
| C20:0 | 0.432 | 0.396 | 0.332 | 0.332 | 0.343 | 0.208 | 0.168 | 0.196 | 0.200 | 0.167 |

[1]Notation indicates carbon chain length followed by number of double bonds. Positional isomers for C14:1, C16:1, and C18:1 FA species are indicated by “a”, b”, and “c”.
[2]Each replicate is comprised of homogenate from 5 flies.
[3]Raw abundance counts are normalized to the abundance of a spiked standard (pentadecanoic acid).

**Supplemental Table 6.** Fatty acid (FA) analysis of male testes in control and antibiotic and antifungal (AB+AF)-treated males for each replicate.

| | Control[3] | | | | | AB+AF[3] | | | | |
|---|---|---|---|---|---|---|---|---|---|---|
| **FA[1,2]** | **1** | **2** | **3** | **4** | **5** | **1** | **2** | **3** | **4** | **5** |
| C10:0 | 0.029 | 0.027 | 0.538 | 0.033 | 0.017 | 0.184 | 0.055 | 0.136 | 0.151 | 0.157 |
| C12:0 | 2.170 | 1.713 | 0.731 | 2.627 | 2.401 | 17.762 | 9.845 | 15.572 | 17.372 | 17.656 |
| C13:0 | 0.000 | 0.012 | 0.000 | 0.000 | 0.000 | 0.035 | 0.023 | 0.047 | 0.024 | 0.049 |
| C14:1a | 0.209 | 0.236 | 0.885 | 0.283 | 0.365 | 0.221 | 0.148 | 0.240 | 0.248 | 0.296 |
| C14:1b | 0.100 | 0.138 | 0.885 | 0.168 | 0.227 | 0.220 | 0.150 | 0.188 | 0.240 | 0.241 |
| C14:1c | 0.332 | 0.324 | 0.885 | 0.417 | 0.399 | 3.646 | 2.381 | 3.547 | 4.000 | 3.688 |
| C14:0 | 3.092 | 2.652 | 0.923 | 3.266 | 2.971 | 16.230 | 12.033 | 15.208 | 18.271 | 17.586 |
| C16:1a | 0.064 | 0.079 | 1.038 | 0.083 | 0.087 | 0.080 | 0.030 | 0.117 | 0.054 | 0.081 |
| C16:1b | 1.545 | 1.520 | 1.038 | 1.638 | 1.622 | 3.769 | 2.802 | 3.704 | 4.355 | 3.870 |
| C16:0 | 7.583 | 7.470 | 1.077 | 7.124 | 6.858 | 9.437 | 6.684 | 10.126 | 10.165 | 11.987 |
| C17:0 | 0.037 | 0.056 | 1.154 | 0.049 | 0.052 | 0.000 | 0.000 | 0.019 | 0.018 | 0.024 |
| C18:2 | 0.234 | 0.248 | 1.192 | 0.232 | 0.200 | 0.182 | 0.096 | 0.198 | 0.171 | 0.219 |
| C18:1a | 2.685 | 2.623 | 1.192 | 2.492 | 2.512 | 2.504 | 1.488 | 2.377 | 2.420 | 2.630 |
| C18:1b | 0.217 | 0.202 | 1.192 | 0.200 | 0.192 | 0.000 | 0.055 | 0.093 | 0.085 | 0.090 |
| C18:0 | 1.138 | 1.119 | 1.231 | 1.047 | 1.203 | 1.124 | 0.752 | 0.916 | 1.097 | 1.159 |
| C20:0 | 0.138 | 0.149 | 1.346 | 0.170 | 0.193 | 0.154 | 0.117 | 0.139 | 0.189 | 0.221 |
| C22:0 | 0.019 | 0.034 | 1.462 | 0.032 | 0.041 | 0.023 | 0.022 | 0.024 | 0.040 | 0.050 |

[1]Notation indicates carbon chain length followed by number of double bonds. Positional isomers for C14:1, C16:1, and C18:1 FA species are indicated by “a”, b”, and “c”.
[2]Each replicate is comprised of homogenate from 5 flies.
[3]Raw abundance counts are normalized to the abundance of a spiked standard (pentadecanoic acid).

**Supplemental Table 7.** Fatty acid (FA) analysis of male testes in control and antibiotic (AB)-treated males for each replicate.

| | Control[3] | | | | AB[3] | | | | |
|---|---|---|---|---|---|---|---|---|---|
| **FA[1,2]** | **1** | **2** | **3** | **4** | **1** | **2** | **3** | **4** | **5** |
| C10:0 | 0.000 | 0.005 | 0.000 | 0.000 | 0.002 | 0.000 | 0.000 | 0.001 | 0.001 |
| C12:0 | 0.024 | 0.050 | 0.036 | 0.146 | 0.011 | 0.021 | 0.027 | 0.013 | 0.021 |
| c14:1b | 0.000 | 0.000 | 0.000 | 0.000 | 0.001 | 0.000 | 0.000 | 0.000 | 0.001 |
| c14:1c | 0.000 | 0.004 | 0.000 | 0.014 | 0.001 | 0.000 | 0.005 | 0.001 | 0.005 |
| C14:0 | 0.050 | 0.085 | 0.065 | 0.223 | 0.021 | 0.045 | 0.066 | 0.025 | 0.043 |
| C16:1a | 1.000 | 1.000 | 1.000 | 1.000 | 1.000 | 1.000 | 1.000 | 1.000 | 1.000 |
| C16:1b | 0.000 | 0.000 | 0.000 | 0.000 | 0.001 | 0.000 | 0.000 | 0.001 | 0.002 |
| C16:0 | 0.006 | 0.033 | 0.013 | 0.024 | 0.014 | 0.033 | 0.036 | 0.017 | 0.027 |
| C17:0 | 0.777 | 1.025 | 1.063 | 1.115 | 0.921 | 0.698 | 0.594 | 0.634 | 0.633 |
| C18:2 | 0.009 | 0.014 | 0.010 | 0.010 | 0.009 | 0.008 | 0.007 | 0.008 | 0.008 |
| C18:1a | 0.000 | 0.000 | 0.000 | 0.000 | 0.001 | 0.005 | 0.004 | 0.002 | 0.002 |
| C18:1b | 0.014 | 0.047 | 0.030 | 0.032 | 0.036 | 0.098 | 0.083 | 0.049 | 0.044 |
| C18:0 | 0.000 | 0.012 | 0.017 | 0.019 | 0.003 | 0.000 | 0.000 | 0.003 | 0.008 |
| C20:0 | 0.361 | 0.469 | 0.485 | 0.406 | 0.457 | 0.253 | 0.213 | 0.257 | 0.241 |
| C22:0 | 0.008 | 0.016 | 0.009 | 0.008 | 0.006 | 0.008 | 0.007 | 0.005 | 0.010 |

[1]Notation indicates carbon chain length followed by number of double bonds. Positional isomers for C14:1, C16:1, and C18:1 FA species are indicated by “a”, b”, and “c”.
[2]Each replicate is comprised of homogenate from 5 flies.
[3]Raw abundance counts are normalized to the abundance of a spiked standard (pentadecanoic acid).

**Supplemental Table 8.** Fatty acid (FA) analysis of male testes from control and antifungal (AF)-treated males for each replicate.

| | Control[3] | | | | | AF[3] | | | | |
|---|---|---|---|---|---|---|---|---|---|---|
| **FA[1,2]** | **1** | **2** | **3** | **4** | **5** | **1** | **2** | **3** | **4** | **5** |
| C10:0 | 0.004 | 0.011 | 0.003 | 0.010 | 0.004 | 0.007 | 0.011 | 0.012 | 0.008 | 0.002 |
| C12:0 | 0.036 | 0.284 | 0.354 | 0.282 | 0.036 | 0.052 | 0.046 | 0.049 | 0.040 | 0.055 |
| C14:1a | 0.000 | 0.000 | 0.010 | 0.004 | 0.000 | 0.000 | 0.000 | 0.000 | 0.000 | 0.000 |
| C14:1b | 0.000 | 0.000 | 0.010 | 0.005 | 0.000 | 0.000 | 0.000 | 0.000 | 0.000 | 0.000 |
| C14:1c | 0.003 | 0.030 | 0.062 | 0.024 | 0.003 | 0.000 | 0.000 | 0.000 | 0.000 | 0.004 |
| C14:0 | 0.038 | 0.238 | 0.350 | 0.248 | 0.038 | 0.079 | 0.062 | 0.060 | 0.047 | 0.059 |
| C16:1b | 0.009 | 0.024 | 0.068 | 0.024 | 0.009 | 0.011 | 0.012 | 0.007 | 0.005 | 0.006 |
| C16:1c | 0.000 | 0.000 | 0.008 | 0.000 | 0.000 | 0.000 | 0.000 | 0.000 | 0.000 | 0.002 |
| C16:0 | 0.678 | 0.987 | 0.921 | 0.965 | 0.678 | 1.112 | 1.367 | 1.144 | 1.302 | 0.714 |
| C17:0 | 0.008 | 0.023 | 0.007 | 0.014 | 0.008 | 0.026 | 0.024 | 0.025 | 0.015 | 0.006 |
| C18:1a | 0.011 | 0.025 | 0.045 | 0.014 | 0.011 | 0.048 | 0.065 | 0.037 | 0.011 | 0.007 |
| C18:1b | 0.005 | 0.013 | 0.013 | 0.010 | 0.005 | 0.023 | 0.029 | 0.018 | 0.010 | 0.006 |
| C18:0 | 0.254 | 0.506 | 0.259 | 0.333 | 0.254 | 0.604 | 0.815 | 0.608 | 0.627 | 0.256 |
| C20:0 | 0.004 | 0.025 | 0.005 | 0.014 | 0.004 | 0.030 | 0.034 | 0.025 | 0.015 | 0.003 |
| C22:0 | 0.000 | 0.003 | 0.000 | 0.003 | 0.000 | 0.009 | 0.013 | 0.000 | 0.000 | 0.000 |

[1]Notation indicates carbon chain length followed by number of double bonds. Positional isomers for C14:1, C16:1, and C18:1 FA species are indicated by “a”, b”, and “c”.
[2]Each replicate is comprised of homogenate from 5 flies.
[3]Raw abundance counts are normalized to the abundance of a spiked standard (pentadecanoic acid).

**Supplemental Table 9.** Fatty acid (FA) analysis of male testes in control and antibiotic- and antifungal (AB+AF)-treated males for each replicate.

| | Control[3] | | | | | AB+AF[3] | | | | |
|---|---|---|---|---|---|---|---|---|---|---|
| **FA[1,2]** | **1** | **2** | **3** | **4** | **5** | **1** | **2** | **3** | **4** | **5** |
| C12:0 | 0.013 | 0.012 | 0.011 | 0.010 | 0.015 | 0.265 | 0.035 | 0.183 | 0.118 | 0.110 |
| C14:1a | 0.000 | 0.001 | 0.000 | 0.001 | 0.001 | 0.004 | 0.000 | 0.003 | 0.001 | 0.001 |
| C14:1b | 0.000 | 0.000 | 0.000 | 0.000 | 0.000 | 0.005 | 0.000 | 0.003 | 0.001 | 0.001 |
| C14:1c | 0.001 | 0.002 | 0.002 | 0.001 | 0.001 | 0.058 | 0.009 | 0.039 | 0.016 | 0.021 |
| C14:0 | 0.030 | 0.042 | 0.037 | 0.034 | 0.034 | 0.355 | 0.049 | 0.306 | 0.161 | 0.162 |
| C16:1a | 0.002 | 0.002 | 0.000 | 0.002 | 0.002 | 0.003 | 0.000 | 0.002 | 0.001 | 0.000 |
| C16:1b | 0.018 | 0.029 | 0.021 | 0.022 | 0.021 | 0.090 | 0.019 | 0.076 | 0.023 | 0.030 |
| C16:0 | 0.601 | 0.598 | 0.564 | 0.595 | 0.559 | 0.714 | 0.554 | 0.731 | 0.761 | 0.768 |
| C17:0 | 0.011 | 0.010 | 0.010 | 0.011 | 0.010 | 0.010 | 0.008 | 0.011 | 0.011 | 0.011 |
| C18:2 | 0.002 | 0.003 | 0.002 | 0.003 | 0.003 | 0.007 | 0.001 | 0.002 | 0.001 | 0.002 |
| C18:1a | 0.048 | 0.067 | 0.051 | 0.052 | 0.062 | 0.079 | 0.030 | 0.068 | 0.027 | 0.033 |
| C18:1b | 0.003 | 0.004 | 0.002 | 0.003 | 0.003 | 0.007 | 0.002 | 0.005 | 0.012 | 0.013 |
| C18:0 | 0.378 | 0.334 | 0.357 | 0.336 | 0.316 | 0.324 | 0.302 | 0.396 | 0.418 | 0.442 |
| C20:0 | 0.008 | 0.010 | 0.009 | 0.008 | 0.010 | 0.020 | 0.006 | 0.012 | 0.010 | 0.013 |

[1]Notation indicates carbon chain length followed by number of double bonds. Positional isomers for C14:1, C16:1, and C18:1 FA species are indicated by “a”, b”, and “c”.
[2]Each replicate is comprised of homogenate from 5 flies.
[3]Raw abundance counts are normalized to the abundance of a spiked standard (pentadecanoic acid).
